# Data-driven dissection of the fever effect in autism spectrum disorder

**DOI:** 10.1101/2022.06.16.496362

**Authors:** Efrat Muller, Ido Shalev, Eitan Bachmat, Alal Eran

## Abstract

Some individuals with autism spectrum disorder (ASD) demonstrate marked behavioral improvements during febrile episodes, in what is perhaps the only present-day means of modulating the core ASD phenotype. Understanding the nature of this so-called *fever effect* is therefore essential for leveraging this natural temporary relief of symptoms to a sustained efficacious intervention. Towards this goal, we used machine learning to analyze the rich clinical data of the Simons Simplex Collection, in which 1 out of every 6 children with ASD was reported to improve during febrile episodes, across multiple ASD domains. Reported behavioral improvements during febrile episodes were associated with maternal infection in pregnancy (OR = 1.7, 95% CI = [1.42, 2.03], P = 4.24×10^−4^) and gastrointestinal (GI) dysfunction (OR=1.46, 95% CI = [1.15, 1.81], P = 1.94×10^−3^). Family members of children reported to improve when febrile have an increased prevalence of autoimmune disorders (OR=1.43, 95% CI = [1.23, 1.67], P = 3.0×10^−6^), language disorders (OR=1.63, 95% CI = [1.29, 2.04], P = 2.5×10^−5^), and neuropsychiatric disorders (OR=1.59, 95% CI = [1.34, 1.89], P < 1×10^−6^). Since both GI abnormalities and maternal immune activation have been linked to ASD via proinflammatory cytokines, these results might suggest a possible involvement of immune dysregulation in the fever effect, consistent with findings in mouse models. This work advances our understanding of the fever-responsive ASD subtype and motivates future studies to directly test the link between proinflammatory cytokines and behavioral modifications in individuals with ASD.

**Lay summary:** Some individuals with ASD demonstrate marked behavioral improvements when they have a fever. However, the magnitude, scope, nature, and underlying neurobiological basis of the so-called fever effect remain unknown. This large-scale biomedical data analysis found that children with ASD reported to improve when febrile tend to be those with worse gastrointestinal symptoms and whose mothers reported an infection during pregnancy. These and other findings point to the involvement of proinflammatory cytokines in this phenomenon.

## INTRODUCTION

During the first half of the twentieth century, artificial fever therapy (pyrotherapy) was widely used to treat an array of neuropsychiatric disorders (Epstein, 1936; Raju, 2006). In the second half of the century, caregivers and parents of children with autism have anecdotally reported marked behavioral improvements during febrile episodes (Cotterill, 1985; Sullivan, 1980). Today, ASD remains one of the most debilitating childhood disorders: It affects 2.27% of the US population (Maenner et al., 2021), and requires chronic management (Harrington & Allen, 2014). Understanding the so-called fever effect in ASD could lead to precision approaches to therapy, based on its underlying neurobiological mechanisms.

Toward this goal, a prospective study of behavioral changes associated with fever in children with ASD was the first to document fewer aberrant behaviors in 30 children with ASD during febrile episodes as compared to 30 matched afebrile children (Curran et al., 2007). More recently, a large-scale investigation of parental reports in the Simons Simplex Collection (SSC) found that one in six children with ASD reportedly improved when he or she had a fever, across multiple ASD domains, including temper, communication, social interaction, repetitive behavior, and cognition (Grzadzinski, Lord, Sanders, Werling, & Bal, 2017). Children reported to improve when febrile tend to have more severe ASD symptoms, such as more repetitive behaviors, and lower non-verbal cognitive skills. However, that important study focused on a selected set of variables from the rich SSC dataset, mostly those related to the child’s behavior and demographics. Additional characteristics of fever responsive children remain to be identified. A comprehensive profiling of these children and their families could shed light on the neurobiological mechanisms underlying the fever effect.

Recent studies in animal models of ASD have shown that interleukin-17a (IL-17a), a pro-inflammatory cytokine often secreted during fever, but not fever per se, rescues social deficits in autistic mice by binding its receptor (IL-17Ra) on neurons of the primary somatosensory cortex dysgranular zone (S1DZ). Importantly, IL-17a produced peripherally as part of a typical immune insult was shown to restore sociability in mouse models of ASD caused by maternal immune activation (MIA), but not in other monogenic models of ASD (Reed et al., 2020). The authors attribute the increased exposure of MIA mice to IL-17a during fetal development to increased neuronal sensitivity for IL-17a after birth (“Gloria Choi,” 2022). Accordingly, direct delivery of IL-17a to the S1DZ of monogenic models of ASD resulted in the same behavioral improvements, as measured by the three-chamber social approach assay and the reciprocal social interaction assay (Reed et al., 2020). Thus the ability of S1DZ neurons to sense IL-17a might distinguish animals whose sociability improves during systemic inflammation from those who do not.

IL-17a signaling was also shown to mediate gastrointestinal inflammation in MIA models of ASD, and overall prime the mouse immune system via epigenetic alterations of naïve CD4+T cells (E. Kim et al., 2022). This ancient cytokine was also shown to act as a neuromodulator in C. elegans, regulating worm behavior (Chen et al., 2017; Flynn et al., 2020). Yet, if and how these findings may translate to humans remains to be elucidated. Immune dysregulation is a well-established etiology for a subset of individuals with ASD (Croen et al., 2019; Ramaswami et al., 2020; Robinson-Agramonte et al., 2022), and increasing levels of pro-inflammatory cytokines have been associated with GI dysfunction in ASD (Ashwood & Wakefield, 2006; Jyonouchi, Geng, Streck, & Toruner, 2011; Samsam, Ahangari, & Naser, 2014), as well as with ASD severity (Ashwood et al., 2011; Masi et al., 2017). IL-17a receptors are expressed throughout the developing and adult human brain (Fujitani, Miyajima, Otani, & Liu, 2022), and plasma IL-17a levels were shown to correlate with cognitive function in individuals with Parkinson’s disease (Green, Khosousi, & Svenningsson, 2019) and multiple sclerosis (Trenova et al., 2018).

Identifying the clinical characteristics of individuals with ASD reported to improve when febrile will advance our understanding of the neurobiological mechanisms underlying behavioral modifications in ASD. We therefore used various machine learning approaches to dissect the rich phenotypic data of the SSC (Fischbach & Lord, 2010) and comparatively characterize the fever responsive subgroup of individuals with ASD, in an unbiased, data-driven fashion.

## METHODS

### Patients and Datasets Used

We mined the SSC data repository (version 15), which includes comprehensive clinical, behavioral, genomic, and demographic data from 2,951 families with one child diagnosed with ASD (Fischbach & Lord, 2010). Among the SSC inclusion criteria are proband age between 4 and 17.9 years, Autism Diagnostic Interview–Revised (ADI-R) and Autism Diagnostic Observation Schedule (ADOS) based diagnoses, and minimal nonverbal cognitive abilities as measured by the Differential Ability Scales-II (DAS-II), Wechsler Intelligence Scale for Children (WISC-IV), or Wechsler Abbreviated Scale of Intelligence (WASI). Exclusion criteria included pregnancy or birth complications, nutritional or psychological deprivation, and family members with ASD, mental retardation, or schizophrenia. Informed consent was obtained and approved by local Institutional Review Boards at each of the 12 data collection sites included in the SSC (https://www.sfari.org/resource/simons-simplex-collection-sites/). Consequently, the Boston Children’s Hospital Institutional Review Board has determined that this particular research qualifies as exempt from the requirements of human subjects protection regulations.

### Data Analysis

#### Feature selection

With a goal of pointing to likely underlying neurobiological mechanisms, we focused on medical variables of the SSC Phenotype Dataset, as well as those considered core and commonly used. In all, 404 variables were selected from 29 SSC tables based on these criteria (**Supplementary Table 1**). Of these, 185 sparse and/or closely related variables were aggregated into 29 summary statistics (**Supplementary Table 2)**. Then 44 variables were removed due to having > 30% missing values, and 20 others were removed for having a constant value across the entire cohort. **Supplementary Figure 1** depicts this feature selection process.

**Figure 1.**
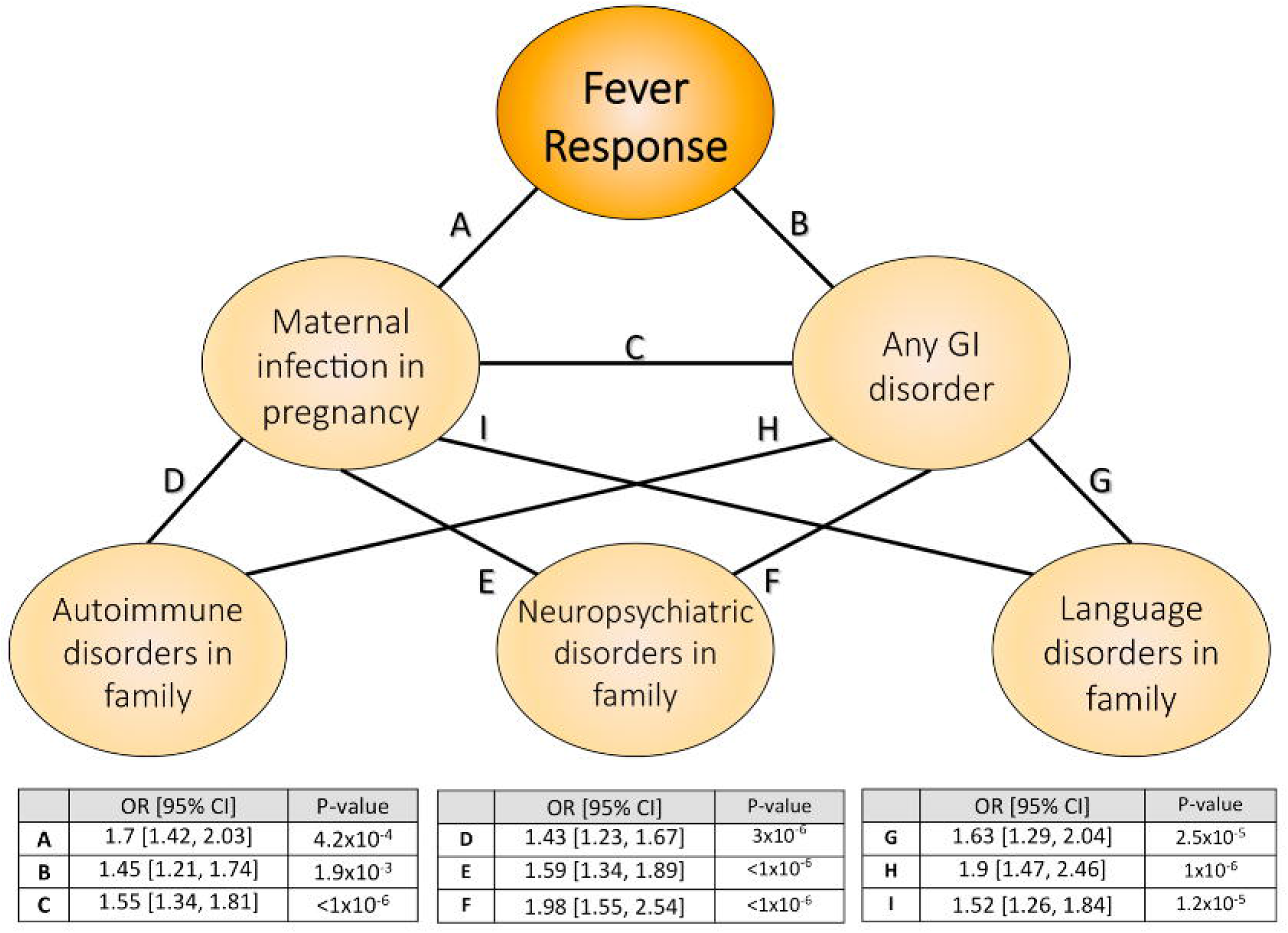
Relationships among domains of improvement during febrile episodes among 2,253 SSC probands. Pairwise overlap in ASD domains of improvement, visualized by Venn diagrams noting the number of individuals reported to improve in either and both domains. ORs and their P values quantify the strength and significance of domain association, respectively. *Significant associations (FDR< 0.05).

#### Continuous variable clustering

Pairwise correlations were calculated between 82 continuous variables. Variables were represented as nodes and edges joined two nodes if their R^2^ ≥ 0.7. Clusters were defined as connected components in this graph. A total of 11 multivariate clusters and 31 single feature clusters were identified (**Supplementary Figure 2** and **Supplementary Table 3**).

**Figure 2.**
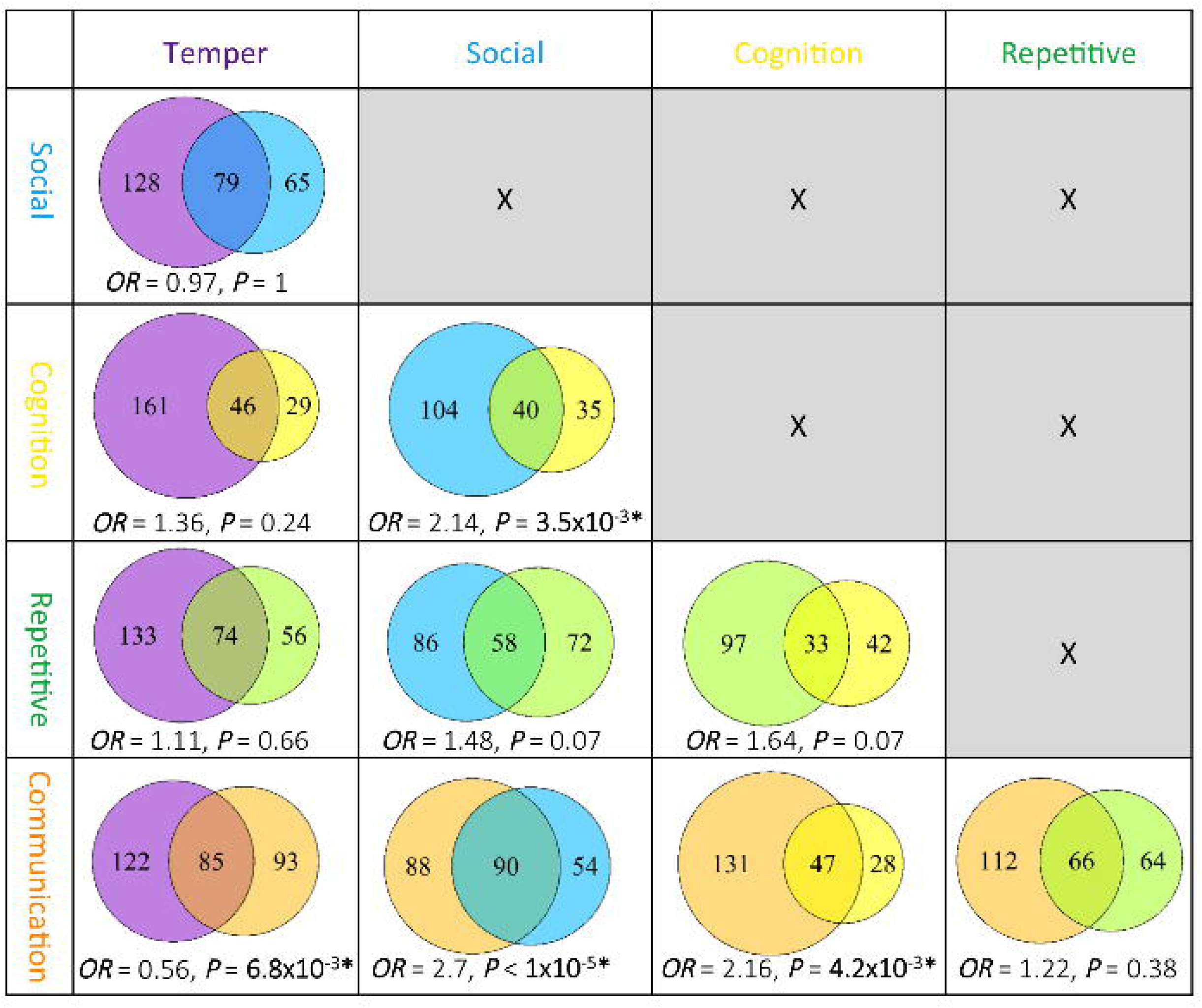
Predictors of fever response. (**A**) Maternal infection in pregnancy is strongly associated with fever response in the proband (OR = 1.7, 95% CI = [1.42, 2.03], P = 4.24×10^−4^). The plot shows the percent of individuals reported to improve when febrile out of all those with answers to questions about maternal infections in pregnancy. (**B**) Fever responders tend to be children with ASD and more severe GI comorbidities (Wilcoxon P = 8.81×10^−18^).

#### Categorical variable clustering

Pairwise Fisher’s exact tests were calculated between 244 categorical variables, using Monte Carlo simulations with B=1×10^7^ where needed. Variables were represented as nodes and edges joined two nodes if their Fisher’s exact association P-value was ≤ 9×10^−6^. This specific P-value threshold was selected as the Benjamini-Hochberg corrected P-value at the jump point in the number of clusters as a function of the P-value (**Supplementary Figure 3A**). Clusters were defined as communities of information flow (Rosvall & Bergstrom, 2008) in this variable association graph. A total of 15 multivariate clusters and 83 single feature clusters were identified using R’s igraph package (Csardi & Nepusz, 2006) (**Supplementary Figure 3B** and **Supplementary Table 4**).

**Figure 3.**
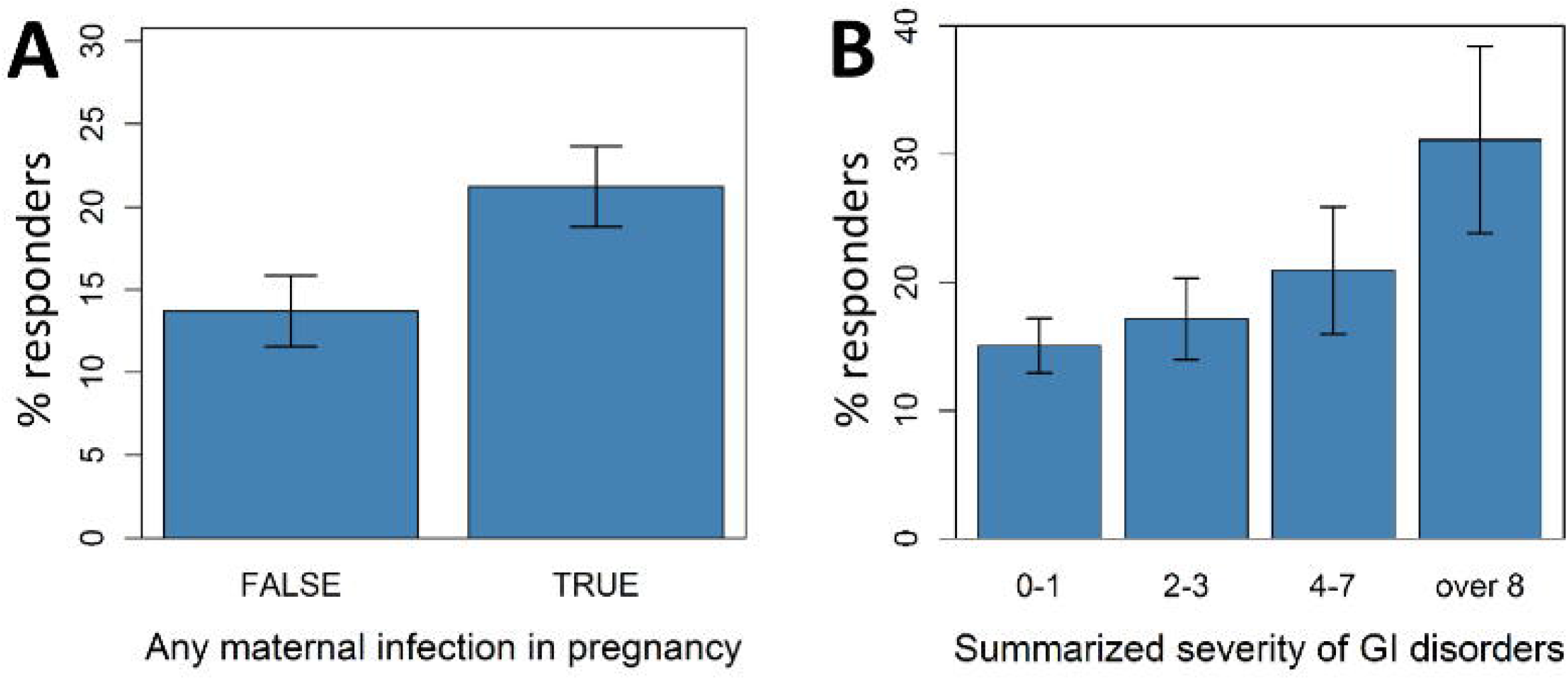
Network structure of the clinical characteristics of children with ASD responding to fever. Shown are clinical features directly and indirectly associated with fever response, their ORs, 95% CIs, and association P values.

#### Characterizing the population of fever responders

Variable clusters described above were tested against a binary target of parental response to whether their child seems to show any improvement in symptoms of autism when s/he has a fever? Children whose parents answered “yes” were labeled as fever responders, those whose parents replied “no” were labeled non-responders, and those whose parents were unsure were ignored. Mann-Whitney U tests were used to assess differences in continuous clusters between responders and non-responders, while Fisher’s exact tests were used for inter-group comparisons of categorical clusters. We used bootstrap validation to stabilize the analysis by repeatedly resampling (n=10,000) equally sized populations from both groups and calculating bootstrapped statistics. Throughout the manuscript, we report median P values and odds ratios (ORs), as well as 95% confidence intervals (CIs) of the bootstrap samples.

#### Domains of improvement analysis

For responders, parental reports included a yes/no answer to observing behavioral changes in the following ASD domains: cognition, social skills, communication, temper, and repetitive behavior. Fisher’s exact tests were used to examine dependencies between domains.

#### Network structure of fever response characteristics

To identify clinical features indirectly associated with fever response, the analysis described under “Characterizing the population of responders” was repeated for the strongest fever response predictors. Specifically, the target variable was changed from fever response to (A) maternal infection in pregnancy, and (B) GI dysfunction. The most associated features were then joined to a network based on their odds ratios.

#### Multivariate predictive power assessment

Continuous and categorical variable clusters found to be related to fever response were integrated into a multivariate classifier using forward feature selection in a support vector machine (SVM), random forest (RF), and generalized linear model (GLM). A subset of 1,340 individuals (235 responders and 1,105 non-responders) that had complete data in all 20 significant variable clusters was included in this analysis. The relative frequency of responders in this sample is consistent with that of the entire SSC. The R packages kernlab (Karatzoglou, Smola, Hornik, & Zeileis, 2004), randomForest (Liaw & Wiener, 2002), and stats (R Core Team, 2014) were used for model fitting. Repeated (n=100) 10-fold cross validation was used to assess the models’ area under the receiver operator characteristics curve (AUC).

#### Assessing the relative importance of maternal awareness

We examined the importance of maternal social cognition in reporting a proband’s fever response relative to other variables in the multivariate classifier described above. A GLM and RF were used to rank variable importance and compare the predictive power of an integrative model fit with and without the maternal social cognition variable. The R packages stats (R Core Team, 2014), randomForest (Liaw & Wiener, 2002), and ROCR (Sing, Sander, Beerenwinkel, & Lengauer, 2005) were used for GLM training, RF modeling, and calculating the area under the receiver operator characteristics curve (AUC), respectively.

#### Multiple testing correction

All P values were corrected for multiple comparisons using the Benjamini-Hochberg procedure (Benjamini & Hochberg, 1995). The false discovery rate (FDR) of this study was controlled at α = 0.05.

## RESULTS

Considering 404 relevant clinical characteristics of the SSC Phenotype Dataset (**Supplementary Table 1** and **Methods**), we identified 17 clusters of continuous variables significantly correlated with fever response (**Supplementary Table 3**), and three clusters of categorical variables significantly associated with fever response (**Supplementary Table 4**). We refer to each cluster with the name of its centroid variable. The most salient differences between fever responders and non-responders were behavioral and cognitive, all previously reported by Grzadzinski et al. in their pioneering analysis of the SSC phenotypic data (**Supplementary Table 3** and Grzadzinski et al., 2017). Here we focus on presenting findings of our unsupervised data-driven analyses that were not previously reported.

### During febrile episodes, 1 in every 6 children with ASD is reported to demonstrate behavioral improvements across associated ASD domains

In accordance with findings first described by Grzadzinski et al. (2017), of 2,253 children with ASD and parental reports of behavior when febrile, 377 (16.7%, 95% CI = [15.2%, 18.3%]) were observed to exhibit improvements in ASD symptoms. Parents of 1,767 probands (78.4%, 95% CI = [76.6%, 80%]) reported no such improvement, and parents of the remaining 109 (4.8%, 95% CI = [4%, 5.8%]) were unsure of their child’s behavioral changes while febrile. First, to understand the nature of the reported fever response, we examined the relationships among ASD domains of improvement (**Figure 1**). Dependencies were detected between improvement in cognition and social behavior (OR = 2.14, 95% CI= [1.24, 3.71], P = 3.5×10^−3^), improvement in cognition and communication (OR = 2.16, 95% CI= [1.24, 3.79], P = 4.2×10^−3^), and, not surprisingly, improvement in communication and social behavior (OR = 2.7, 95% CI= [1.72, 4.26], P < 1×10^−6^). The latter is consistent with their merging into a single ASD domain in the Diagnostic and Statistical Manual of Mental Disorders, Fifth Edition (DSM-5) (American Psychiatric Association, 2013). Furthermore, we found that probands that improve in temper when febrile were less likely to exhibit communication improvements (OR = 0.56, 95% CI= [0.36, 0.86], P= 6.8×10^−3^). We further compared improvement in any one domain to improvement in the remaining four domains, to test whether any specific domain stands out on its own. Improvement in each domain was strongly associated with improvement in the other domains (P = 1×10^−8^, Fisher’s exact test, **Supplementary Figure 4**), implying that fever responders tend to improve in several ASD domains and not in any specific one.

### Maternal infection in pregnancy is strongly associated with fever response in children with ASD

Mothers of children reported to improve when febrile also tended to report having had an infection during pregnancy, including herpes, influenza, and enterovirus infections (**Figure 2A**, OR = 1.7, 95% CI = [1.42, 2.03], P = 4.24×10^−4^). Specifically, maternal infection in pregnancy is associated with improvement in repetitive behavior (OR=3.15, 95% CI = [2.14-4.96], P < 1.00×10^−5^), improvement in social interaction (OR = 2.32, 95% CI = [1.63,3.45], P = 3.03×10^−6^), and improvement in communication skills (OR = 1.84, 95% CI = [1.34,2.56], P = 1.68×10^−4^).

### Fever response is strongly related to GI dysfunction in children with ASD

Children reported to improve during febrile episodes were more likely to be reported as suffering from various GI dysfunctions, including diarrhea, bloating, and severe abdominal pain (**Supplementary Figure 5**, OR=1.46, 95% CI = [1.15, 1.81], P = 1.94×10^−3^). The more severe the overall GI symptoms, the greater the chance for observed behavioral improvement during febrile episodes (**Figure 2B**, Wilcoxon P = 8.81×10^−18^).

### Network analysis identifies relationships between fever response, maternal infection in pregnancy, and GI dysfunction

To clinically characterize the ASD subtype of fever responders, we also examined clinical features indirectly associated with fever response. This analysis revealed that autoimmune disorders in the family, neuropsychiatric disorders in the family, and language disorders in the family are all characteristics of fever response, linked via GI dysfunction and maternal infection during pregnancy (**Figure 3**).

### Maternal social cognition confounds fever response characterization

Maternal social cognition scores, as measured by the SRS, were lower for mothers of children reported to improve when febrile, reflecting a greater maternal ability to interpret social cues once they are picked up (**Supplementary Figure 6A**, P=3.94×10^−3^). No relationship was detected between paternal social cognition and a proband’s fever response (**Supplementary Figure 6B**, P = 0.17). Since SSC mothers tend to be those responsible for most parental reports, this finding suggests that maternal social abilities represent a confounder in this analysis, which future study designs should minimize.

### Multivariate analysis demonstrates that further fever effect determinants remain to be discovered

We used multivariate modeling to understand the overall and relative contributions of the identified fever effect determinants. Forward feature selection was first applied to identify the optimal combination of clinical characteristics most informative of fever response (**Supplementary Figure 7**). Using repeated 10x cross validation, we found that logistic regression with nine variables predicted fever response with a mean AUC of 0.696 (0.056) (**Supplementary Figure 7**). Variable ranking identified the most informative determinants to be verbal IQ, ADI-R verbal communication score, and ADI-R repetitive and restricted behavior (RRB) score, all previously reported by Grzadzinski et al. (**Supplementary Figure 8**). We used two complementary approaches to assess the impact of the maternal SRS social cognition confounder. In the first, we excluded this variable from the optimal model and compared the model’s performance with and without the maternal social cognition variable. Compared to the full model’s mean AUC of 0.696 (0.056), the maternal SRS social cognition –null model performed with a mean AUC of 0.678 (0.059), suggesting that although maternal cognition confounds our analysis, its overall contribution to the fever effect characterization is modest (**Supplementary Figure 9**). In the second approach, we matched responders and non-responders based on their maternal SRS social cognition scores, and tested the predictive power of the optimal multivariate model on these subpopulations. The resulting mean AUC of 0.679 (0.061) (**Supplementary Figure 10**) suggests that further fever effect determinants remain to be discovered.

## DISCUSSION

This data-driven analysis of the fever effect in ASD provides a comprehensive characterization of the fever-responsive ASD subtype. The previous study analyzing the same data focused mainly on behavioral and cognitive characteristics of fever responders, highlighting the role of repetitive and restrictive behaviors and lower non-verbal IQ as their most distinctive features (Grzadzinski et al., 2017). The machine learning approaches utilized in our study represent complementary analyses that support and extend their findings, focusing on medical characteristics of probands and their family members. While the non-behavioral differences are more modest than the behavioral ones, our network analysis suggests that they could hold the key to understanding how the fever effect is linked with other neuropsychiatric, autoimmune, and language disorders, and point to likely neurobiological mechanisms.

The strong association of familial characteristics with fever response predictors suggests that fever response in ASD might have a substantial inherited component. Moreover, since each of these features has been linked to pro-inflammatory cytokine dysregulation, namely the interleukin (IL)-17, IL-6, and IL-2 pathways (Choi et al., 2016; Estes & McAllister, 2016; Ferguson et al., 2016; Hsiao, McBride, Chow, Mazmanian, & Patterson, 2012; Hsiao et al., 2013; Jyonouchi, Geng, Ruby, & Zimmerman-Bier, 2005; Jyonouchi et al., 2011; Samsam et al., 2014; Shin Yim et al., 2017), our findings hint to a likely underlying neurobiological mechanism in humans.

Our work is consistent with recent studies in ASD animal models exposed to maternal immune activation (MIA) in utero. Specifically, findings in mice show that MIA causes elevated production of proinflammatory cytokines such as IL-17a (Choi et al., 2016; Kalish et al., 2021; Reed et al., 2020), which cross the placenta and affect mRNA translation in the developing fetal brain, increasing the risk for neurodevelopmental disorders such as ASD (Kalish et al., 2021; Reed et al., 2020). Moreover, IL-17a was shown to alter the maternal gut microbiota, which renders the immune system of the offspring more susceptible to GI inflammatory responses (Eunha Kim et al., 2022; Kim et al., 2017). Importantly, MIA models of ASD show improvement in social behaviors during lipopolysaccharide (LPS)-induced inflammation, mediated by elevated IL-17a production (Reed et al., 2020). Accordingly, our findings provide the first evidence for a subset of human subjects with ASD exposed to maternal immune activation in utero, with comorbid GI dysfunction, and improved behaviors during febrile episodes. Our findings suggest that proinflammatory cytokines might underly the reported beneficial effects of fever in this subgroup.

Moreover, the association of familial autoimmune, neuropsychiatric, and language disorders with this subtype provides further support for the involvement of pro-inflammatory cytokines, since these disorders share a well-documented immune dysregulation etiology (Cerri, Caleo, & Bozzi, 2017; Estes & McAllister, 2016; Hunter & Jones, 2015; Knuesel et al., 2014; Tabarkiewicz, Pogoda, Karczmarczyk, Pozarowski, & Giannopoulos, 2015). One implication of our findings is that proinflammatory cytokines might also have beneficial effects in these disorders, consistent with findings in MIA mice. Indeed, several anecdotal reports describe symptom amelioration in some individuals with attention deficit hyperactivity disorder (ADHD) (Marner), schizophrenia (Zuschlag, Lalich, Short, Hamner, & Kahn, 2016), and bipolar disorder (BP) (Setsaas & Vaaler, 2014) during febrile episodes. Further research should characterize the fever effect in these neurodevelopmental disorders. Though some changes in behavior at times of illness are expected (e.g., higher lethargy, reduced aggressiveness), improvements such as enhanced communication skills, less repetitive behaviors, or cognitive improvements are surprising. For ASD, a common neurodevelopmental disorder with no pharmacological treatment options, this natural remedy represents one of few leads toward therapy.

Our study has several limitations. Its main limitation stems from its source of data, namely retrospective parent reports. A reliance on retrospective memory introduces several biases to the data. While major medical events and demographic features during pregnancy (such as time of pregnancy) are reported accurately, other events, including maternal infection, are typically underestimated (Buka, Goldstein, Spartos, & Tsuang, 2004; Voldsgaard et al., 2002). Parental reports might introduce additional bias, as evidenced by the confounding effect of the mother’s social cognition. Besides that, other parental biases not directly captured in the data might have affected our results. These include the amount of time the parents spend with their child, parental tendency to track symptoms and be alerted to changes, and the frequency of febrile episodes in the child.

Another limitation of this study is that by basing our analyses on a simple “yes/no” target we ignore the degree of improvement, the possibility of an opposite effect (i.e., worse behavior), the fever effect’s dynamics throughout development, and more precise diagnoses reflecting the cause of fever during which improvement was observed. Additionally, other factors may influence a child’s behavior when he or she are sick, including increased parental or caregiver stress, increased attention, changes in sleep patterns, and changes in nutrition.

Follow up prospective studies could resolve these limitations by (A) minimizing parental reporting biases, e.g., with the help of technology, (B) dissecting the immune processes accompanying behavioral changes, (C) incorporating electronic health record data for a more precise characterization of the fever-responsive ASD subtype, and (D) fine-mapping the behavioral domains that do and do not vary in response to immune insults. For example, a smartphone app could be used to track behavioral changes and comorbid conditions in real time.

## CONCLUSIONS

An emerging subtype of fever-responsive ASD may be characterized by GI abnormalities, maternal infection in pregnancy, and familial autoimmune and neuropsychiatric disorders. Our work suggests that the fever effect might be shared across neurodevelopmental disorders, implying that similar analyses would be informative for dissecting ADHD, schizophrenia, and BP. Our findings suggest that future studies of circulating pro-inflammatory cytokine dynamics and their relations to fever and behavioral changes in human subjects could pave the way towards the development of targeted treatment options for this ASD subtype and other immune-mediated neurodevelopmental disorders.

## Acknowledgments

We are grateful to all of the families at the participating Simons Simplex Collection (SSC) sites, as well as the principal investigators (A. Beaudet, R. Bernier, J. Constantino, E. Cook, E. Fombonne, D. Geschwind, R. Goin-Kochel, E. Hanson, D. Grice, A. Klin, D. Ledbetter, C. Lord, C. Martin, D. Martin, R. Maxim, J. Miles, O. Ousley, K. Pelphrey, B. Peterson, J. Piggot, C. Saulnier, M. State, W. Stone, J. Sutcliffe, C. Walsh, Z. Warren, E. Wijsman). We appreciate obtaining access to phenotypic data on SFARI Base. Approved researchers can obtain the SSC population dataset described in this study (https://base.sfari.org/ordering/phenotype/sfari-phenotype/download?code=11) by applying at https://base.sfari.org. We thank members of the Eran lab for fruitful discussions. This work was supported by the Israel Science Foundation (grant 2755/20).

