## Supplementary Information for "Data-driven dissection of the fever effect in autism spectrum disorder"

**This file includes:**

Supplementary Figures 1 to 10

Supplementary Tables 1 to 4

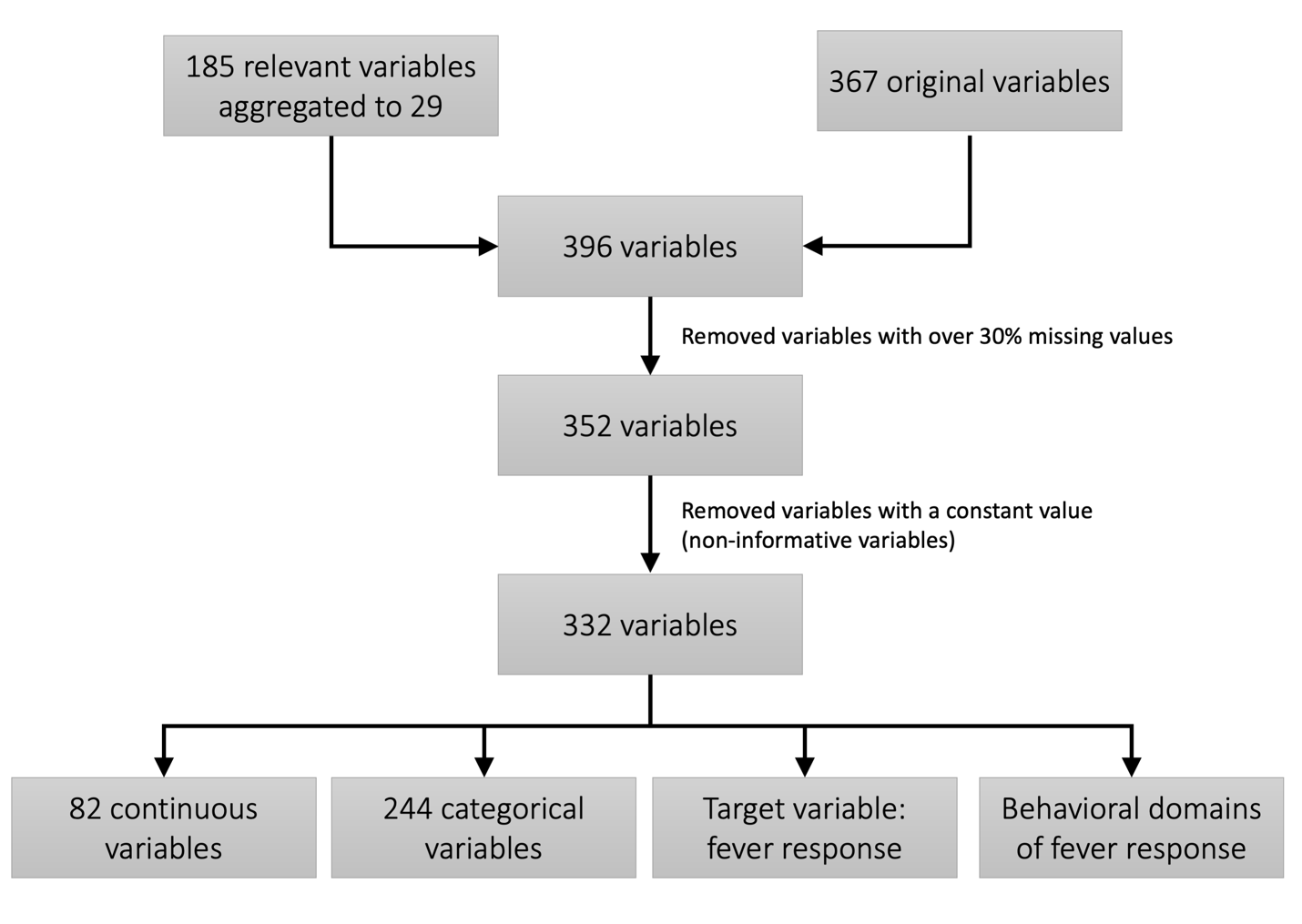

**Supplementary Figure 1** Overview of the feature selection process**.**

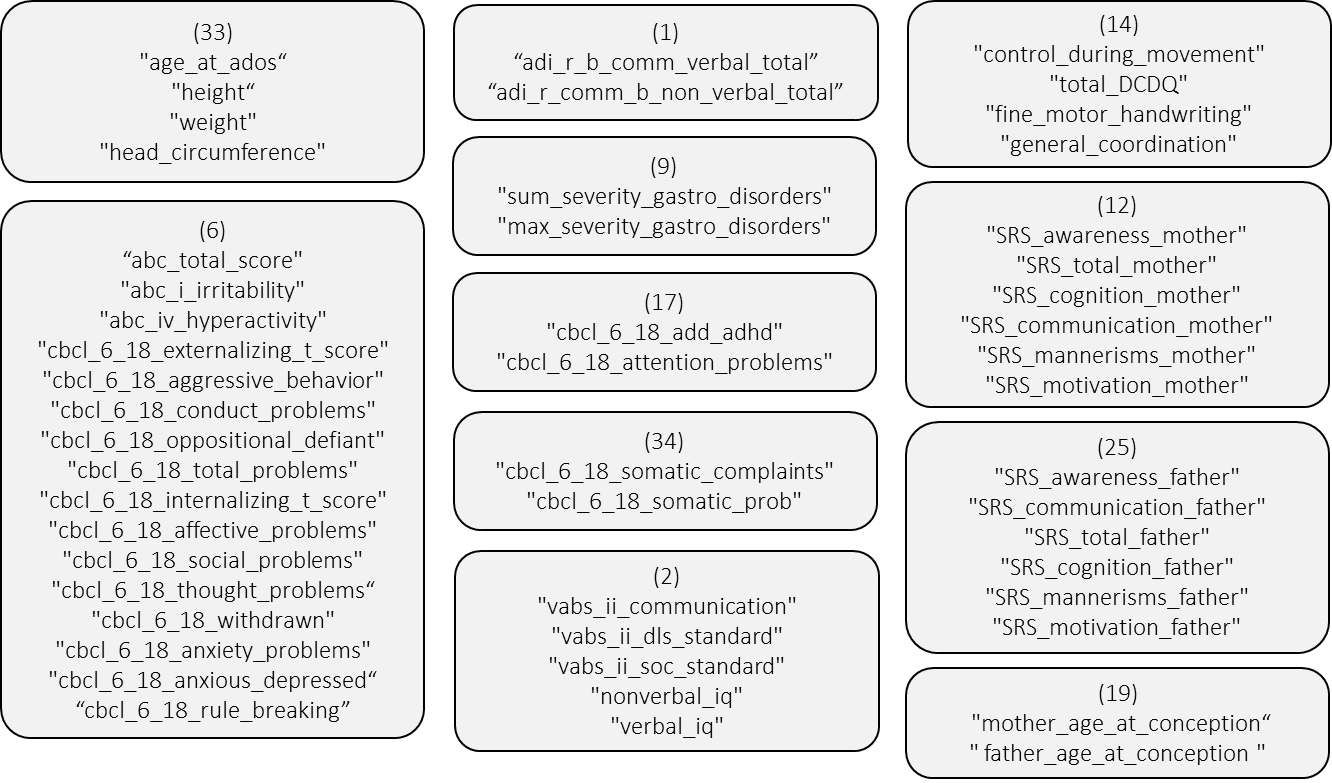

**Supplementary Figure 2** Continuous variable clusters. Shown are 11 multivariate clusters, identified as minimal spanning trees of continuous variables connected by an edge if their pairwise correlation was ≥ 0.7 across the entire cohort. Additionally, 31 clusters of size 1 were identified (**Supplementary** **Table 3**).

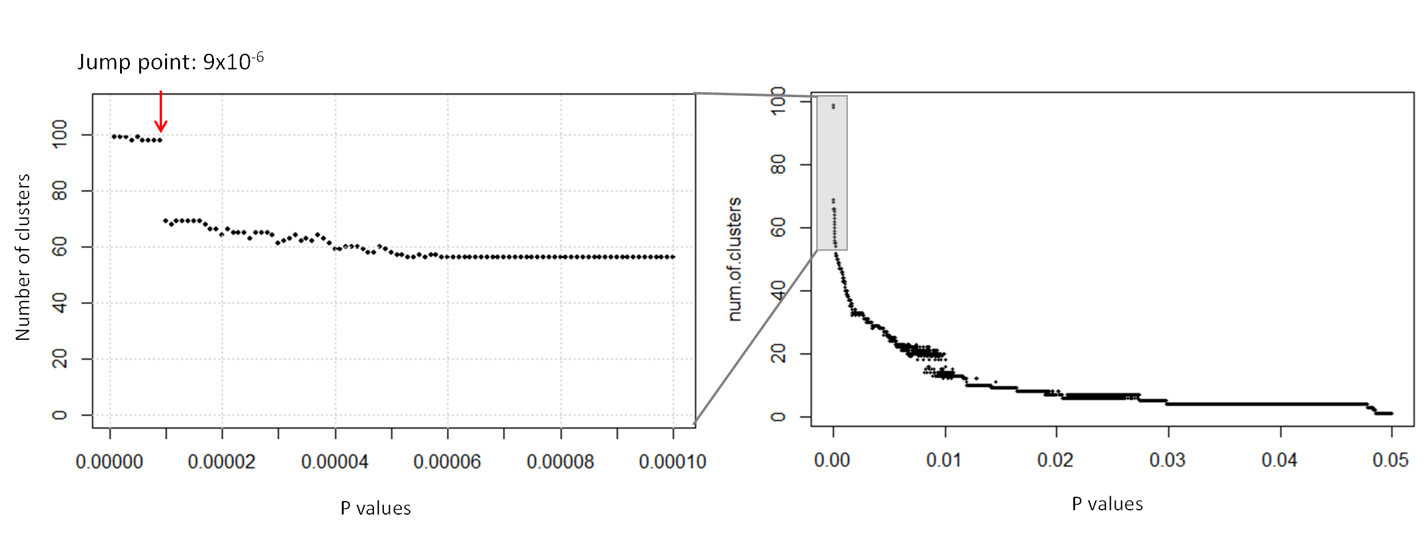
**(A)**

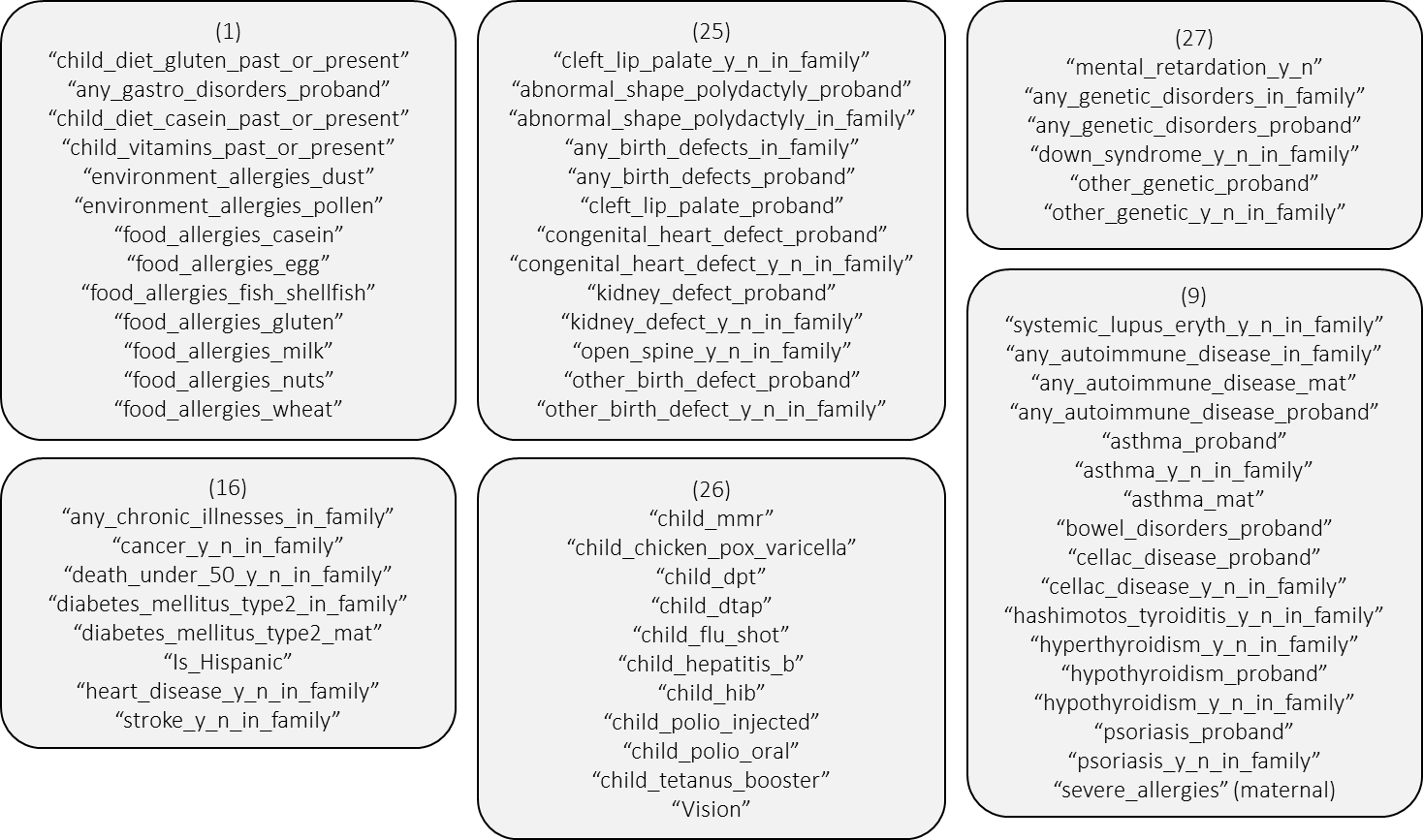
**(B)**

**Supplementary Figure 3** Categorical variable clusters. (**A**) Plotting the number of clusters as a function of pairwise Fisher’s Exact P-values revealed a jump point at P = 9x10-6. This P value was

selected as the cutoff for connecting associated variables into cliques, thereby forming categorial clusters. On the right are the number of clusters as a function of the P values from 0 to 0.05, and on the left is a zoomed in P value range of 0 to 0.005. (**B**) Shown are 6 of 18 multivariate clusters, identified as maximal cliques of categorical variables connected by an edge if their pairwise Fisher’s Exact P value was ≤ 9x10-6. Additionally, 86 clusters of size 1 were identified (Supplementary Table 3).

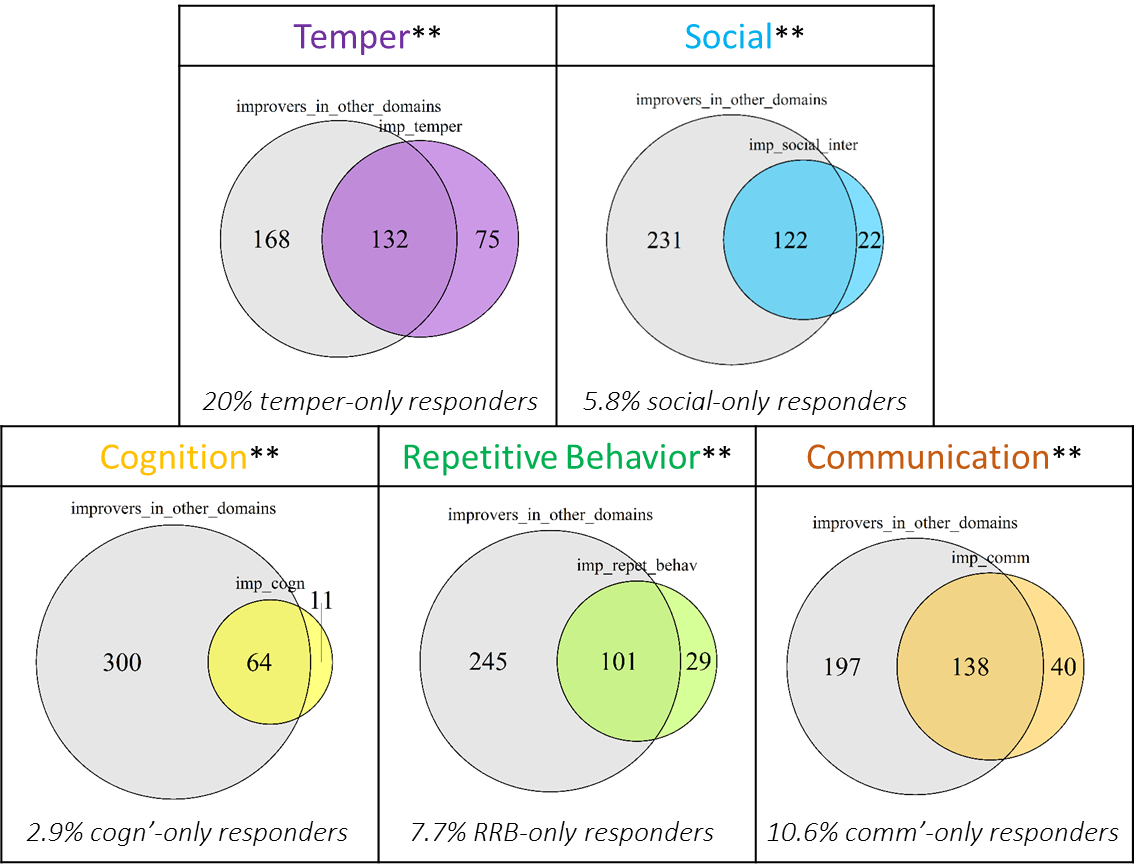

**Supplementary Figure 4** Relationships between improvements across ASD domains during febrile episodes. Venn diagrams illustrate the pairwise overlap between individuals reported to improve in a certain domain as compared to other domains, noting the number of individuals in either and both groups. Below each diagram, the percentage of individuals reported to demonstrate domain-specific improvement is noted (out of all fever responders). **Fisher’s exact test P < 1x10^-6^. Cogn’, cognition; RRB, restricted and repetitive behavior; comm’, communication.

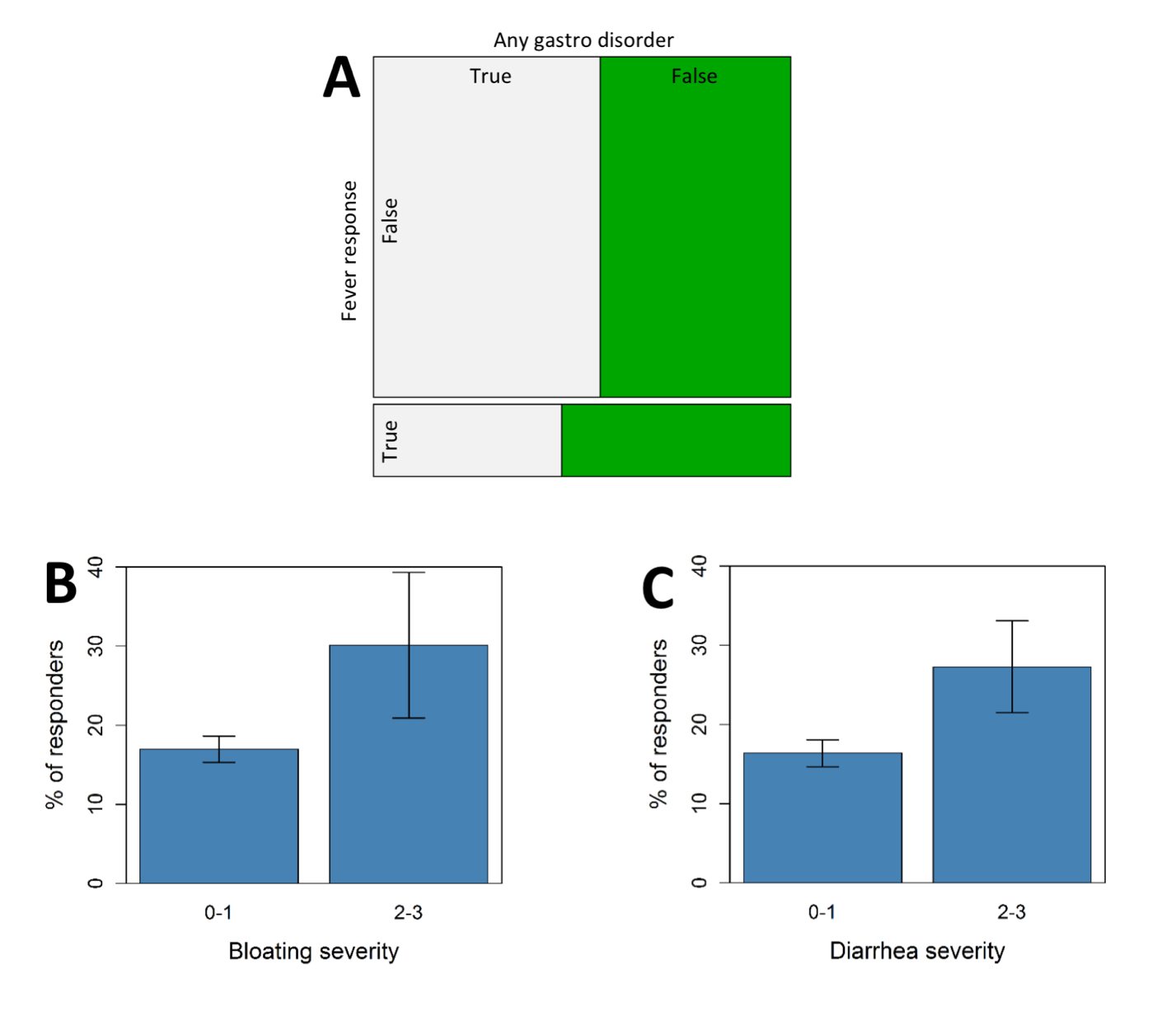

**Supplementary Figure 5** Relationships between gastrointestinal dysfunction and fever response. (**A)** A mosaic plot depicts the association between fever response and any gastrointestinal problem in the proband, as defined in Supplementary Table 2. (OR=1.46, 95% CI = [1.15, 1.81], median bootstrapped Fisher’s P = 1.94x10^-3^]. Specifically, individuals with ASD reported to improve when febrile tend to experience (**B**) more severe bloating (median bootstrapped Wilcoxon P = 1.54x10^-2^), and (**C**) more severe diarrhea (median bootstrapped Wilcoxon P = 2.57x10^-3^). Error bars represent 95% confidence intervals.

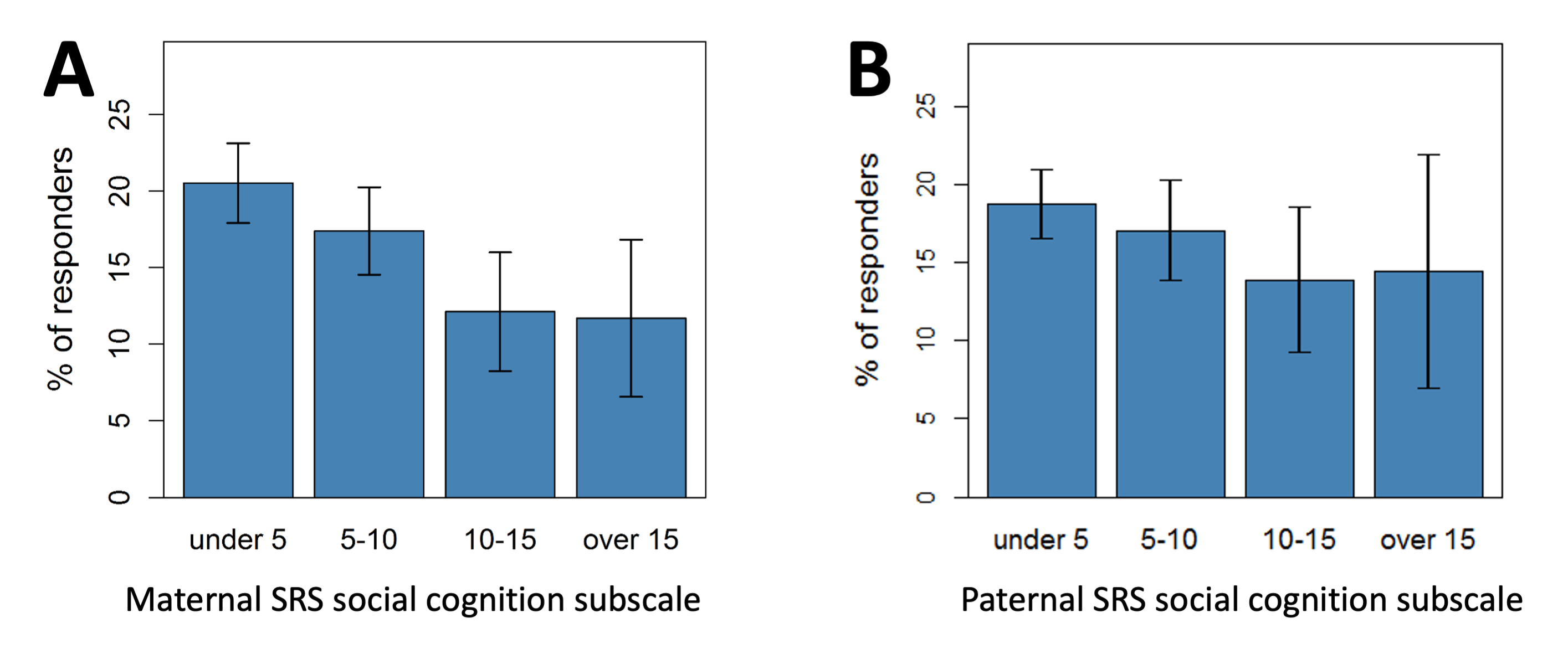

**Supplementary Figure 6** Relationships between parental SRS social cognition scores and fever response. These scores represent a measure of social cognition and not cognitive ability per se. Lower scores reflect a greater ability to interpret social cues. Shown is the percent of SSC probands reported to improve when febrile per (**A**) maternal SRS social cognition scores (Wilcoxon P = 3.94x10^-3^), and (**B**) paternal SRS social cognition scores (Wilcoxon P = 0.17). Error bars represent 95% confidence intervals

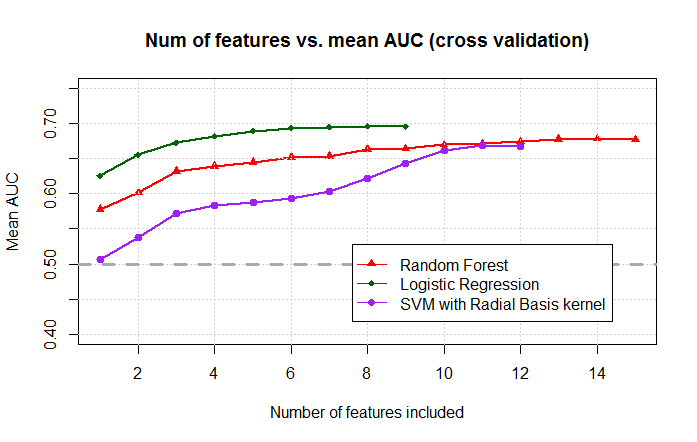
**A**

| Model | Step | Feature Added | AUC mean | AUC SD |
| --- | --- | --- | --- | --- |
| LR | 1 | adi_r_rrb_c_total | 0.626 | 0.063 |
| LR | 2 | verbal_iq | 0.656 | 0.061 |
| LR | 3 | SRS_social_cognition_mother | 0.673 | 0.057 |
| LR | 4 | abc_total_score | 0.682 | 0.059 |
| LR | 5 | any_maternal_infections_in_preg | 0.689 | 0.058 |
| LR | 6 | child_diet_gluten_past_or_present | 0.693 | 0.058 |
| LR | 7 | adi_r_b_comm_verbal_total | 0.695 | 0.058 |
| LR | 8 | abc_iii_stereotypy | 0.696 | 0.058 |
| LR | 9 | abc_v_inappropriate_speech | 0.696 | 0.056 |

**B**

| Model | Step | Feature Added | AUC mean | AUC SD |
| --- | --- | --- | --- | --- |
| SVM | 1 | adi_r_rrb_c_total | 0.507 | 0.034 |
| SVM | 2 | adi_r_soc_a_total | 0.537 | 0.071 |
| SVM | 3 | total_DCDQ | 0.572 | 0.077 |
| SVM | 4 | bloating_specify | 0.584 | 0.066 |
| SVM | 5 | SRS_social_cognition_mother | 0.588 | 0.063 |
| SVM | 6 | nonstandardized_impressions | 0.593 | 0.063 |
| SVM | 7 | abc_total_score | 0.603 | 0.060 |
| SVM | 8 | verbal_iq | 0.622 | 0.060 |
| SVM | 9 | abc_ii_lethargy | 0.643 | 0.061 |
| SVM | 10 | cbcl_6_18_social | 0.662 | 0.058 |
| SVM | 11 | adi_r_b_comm_verbal_total | 0.669 | 0.061 |
| SVM | 12 | child_diet_gluten_past_or_present | 0.667 | 0.060 |

**C**

| Model | Step | Feature Added | AUC mean | AUC SD |
| --- | --- | --- | --- | --- |
| RF | 1 | verbal_iq | 0.578 | 0.059 |
| RF | 2 | abc_iii_stereotypy | 0.602 | 0.064 |
| RF | 3 | any_maternal_infections_in_preg | 0.632 | 0.061 |
| RF | 4 | diarrhea_specify | 0.639 | 0.060 |
| RF | 5 | cbcl_6_18_attention_problems | 0.645 | 0.064 |
| RF | 6 | abc_total_score | 0.652 | 0.057 |
| RF | 7 | adi_r_soc_a_total | 0.654 | 0.061 |
| RF | 8 | SRS_social_cognition_mother | 0.663 | 0.058 |
| RF | 9 | nonstandardized_impressions | 0.664 | 0.057 |
| RF | 10 | abc_v_inappropriate_speech | 0.670 | 0.057 |
| RF | 11 | sum_severity_gastro_disorders | 0.671 | 0.058 |
| RF | 12 | adi_r_rrb_c_total | 0.674 | 0.058 |
| RF | 13 | abc_ii_lethargy | 0.677 | 0.057 |
| RF | 14 | child_diet_gluten_past_or_present | 0.678 | 0.058 |
| RF | 15 | bloating_specify | 0.677 | 0.056 |

**D**

**Supplementary Figure 7** Forward feature selection using logistic regression (LR), support vector machine (SVM), and random forest (RF) models. (**A**) The mean area under the receiver operating characteristics curve (AUC) of 10-fold repeated cross validations according to the number of features included in each model. (**B-D**) The clinical features most predictive of fever response, as identified by a forward feature selection process using (**B**) LR, (**C**) SVM, and (**D**) RF, according to their selection order.

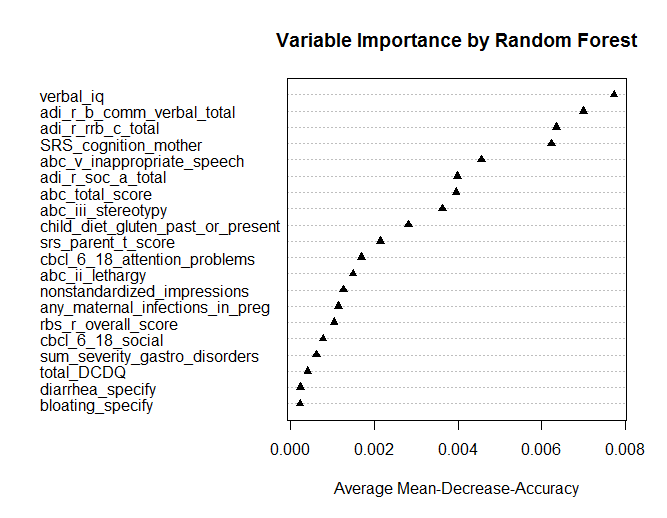

**Supplementary Figure 8** Variable importance according to RF’s mean decrease in accuracy. For each fever response characteristic and each RF tree, the prediction error on the out-of-bag portion of the data was recorded. The same was done after permuting each feature, and the difference between the two were averaged over all trees, and then normalized by the standard deviation of the differences.

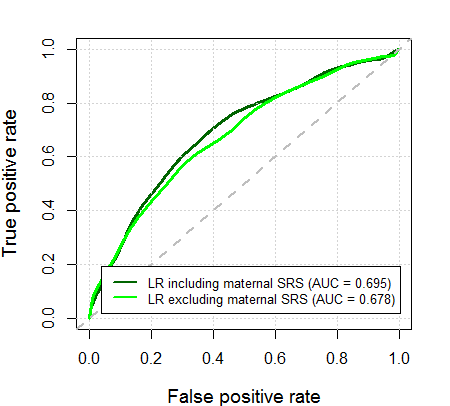

**Supplementary Figure 9** Receiver operating characteristics (ROC) curves of LR models with (dark green) and without (light green) the maternal SRS social cognition feature.

**A**

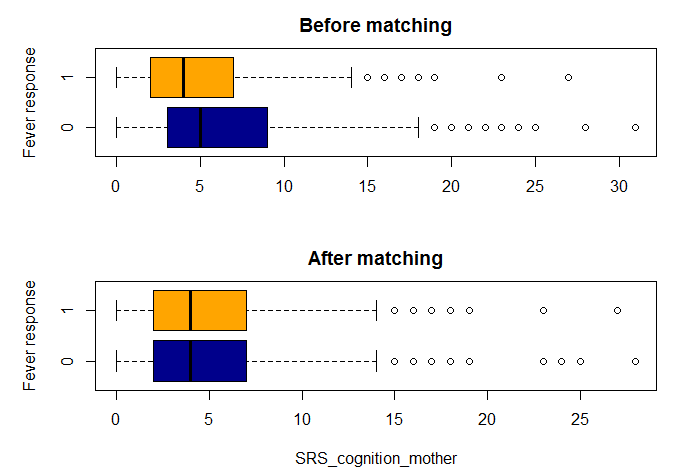

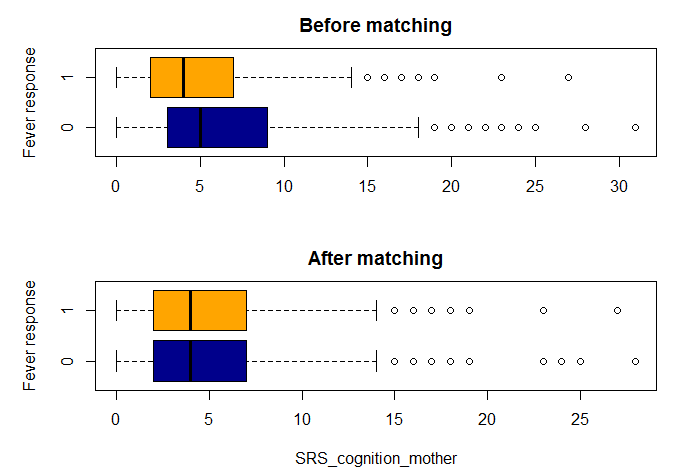

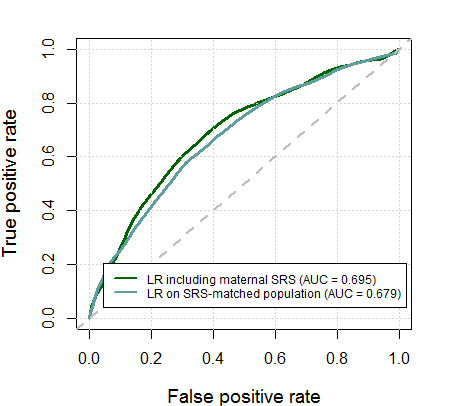
**B**

**Supplementary Figure 10** Assessing the maternal SRS social cognition confounder using matching. (**A**) Boxplots of maternal SRS social cognition scores before and after nearest neighbor matching. Before matching, the optimal LR model was trained on 1,340 individuals (235 responders and 1,105 non-responders). For each of the 235 responders, 4 non-responders with closest maternal SRS social cognition scores were chosen. The 1:4 ratio was chosen to maximize the number of probands included, while maintaining the approximate ratio of responders in the general population. 1,175 individuals (235 responders and 940 non-responders) were included in the maternal SRS-matched cohorts. (**B**) ROC curves of LR models with and without maternal SRS social cognition matching.

**Supplementary Table 1 (separate file).** All variables included in the analysis. The first column denotes each variable’s unique ID, and the second column describes its name. The third column details whether the variable is an aggregated one created by us, or an original SSC variable. The fourth column is relevant only for aggregated variables and includes the aggregation function used to generate them. The fifth column details the original variable name, and the sixth column lists the SSC table from which the variable was taken. The seventh column denotes the original variable type. The eighth to twelfth columns detail the family member to which the variable relates: mother, father, proband, sibling, or other, respectively. Finally, the thirteenth column describes the full variable name, and the fourteenth column includes any notes.

**Supplementary Table 2** Aggregated variables

| No. | New Variable Name | Function Used | Original Variables Used |
| --- | --- | --- | --- |
| 1 | sum_severity_gastro_disorders | SUM | bloating_specify, vomiting_specify, constipation_specify, celiac_disease_gas_ill_specify, diarrhea_specify, unusual_stools_specify, excessive_gas_specify, severe_abdominal_pain_specify, gastroesophageal_reflux_specify, irritable_bowel_specify |
| 2 | max_severity_gastro_disorders | MAX | bloating_specify, vomiting_specify, constipation_specify, celiac_disease_gas_ill_specify, diarrhea_specify, unusual_stools_specify, excessive_gas_specify, severe_abdominal_pain_specify, gastroesophageal_reflux_specify, irritable_bowel_specify |
| 3 | any_gastro_disorders_proband | OR | bloating_specify, vomiting_specify, constipation_specify, celiac_disease_gas_ill_specify,diarrhea_specify, unusual_stools_specify, excessive_gas_specify, severe_abdominal_pain_specify, gastroesophageal_reflux_specify, irritable_bowel_specify |
| 4 | any_autoimmune_disease_proband | OR | bowel_disorders_proband, diabetes_mellitus_type2_proband, hypothyroidism_proband, rheum_arthritis_adult_proband, asthma_proband, diabetes_mellitus_type1_proband, hyperthyroidism_proband, psoriasis_proband, systemic_lupus_eryth_proband, adrenal_insufficiency_proband, hashimotos_tyroiditis_proband, multiple_sclerosis_proband, rheum_arthritis_juvenile_proband, cellac_disease_proband |
| 5 | any_autoimmune_disease_in_family | OR | bowel_disorders_y_n_in_family, diabetes_mellitus_type2_y_n_in_family, hypothyroidism_y_n_in_family, rheum_arthritis_adult_y_n_in_family, asthma_y_n_in_family, diabetes_mellitus_type1_y_n_in_family, hyperthyroidism_y_n_in_family, psoriasis_y_n_in_family, systemic_lupus_eryth_y_n_in_family, adrenal_insufficiency_y_n_in_family, hashimotos_tyroiditis_y_n_in_family, multiple_sclerosis_y_n_in_family, rheum_arthritis_juvenile_y_n_in_family, cellac_disease_y_n_in_family |
| 6 | number_of_distinct_autoimmune  _diseases_in_family | COUNT DISTINCT | bowel_disorders_y_n_in_family, diabetes_mellitus_type2_y_n_in_family, hypothyroidism_y_n_in_family, rheum_arthritis_adult_y_n_in_family, asthma_y_n_in_family, diabetes_mellitus_type1_y_n_in_family, hyperthyroidism_y_n_in_family, psoriasis_y_n_in_family, systemic_lupus_eryth_y_n_in_family, adrenal_insufficiency_y_n_in_family, hashimotos_tyroiditis_y_n_in_family, multiple_sclerosis_y_n_in_family, rheum_arthritis_juvenile_y_n_in_family, cellac_disease_y_n_in_family |
| 7 | any_birth_defects_proband | OR | abnormal_shape_polydactyly_proband, cleft_lip_palate_proband, congenital_heart_defect_proband, other_birth_defect_proband, open_spine_proband, kidney_defect_proband |
| 8 | any_birth_defects_in_family | OR | abnormal_shape_polydactyly_y_n_in_family, cleft_lip_palate_y_n_in_family, congenital_heart_defect_y_n_in_family, other_birth_defect_y_n_in_family, open_spine_y_n_in_family, kidney_defect_y_n_in_family |
| 9 | any_chronic_illnesses_in_family | OR | cancer_y_n_in_family, death_under_50_y_n_in_family, heart_disease_y_n_in_family, stroke_y_n_in_family, other_disorder_illness1_y_n_in_family, other_disorder_illness2_y_n_in_family |
| 10 | any_language_disorders_in_family | OR | communication_disorder_y_n_in_family, expressive_lang_disorder_y_n_in_family, mixed_expressive_disorder_y_n_in_family, pragmatics_lang_disorder_y_n_in_family, stuttering_y_n_in_family, speech_delay_y_n_in_family, reduced_articulation_y_n_in_family |
| 11 | any_language_disorders_proband | OR | communication_disorder_proband, expressive_lang_disorder_proband, mixed_expressive_disorder_proband, pragmatics_lang_disorder_proband, receptive_lang_disorder_proband, stuttering_proband, speech_delay_proband, reduced_articulation_proband |
| 12 | any_genetic_disorders_in_family | OR | angelman_y_n_in_family, cri_du_chat_y_n_in_family, down_syndrome_y_n_in_family, fragile_x_y_n_in_family, phenylketonuria_y_n_in_family, prader_willi_y_n_in_family, rett_syndrome_y_n_in_family, other_genetic_y_n_in_family, smith_lemli_opitz_y_n_in_family |
| 13 | any_genetic_disorders_proband | OR | angelman_proband, cri_du_chat_proband, down_syndrome_proband, fragile_x_proband, phenylketonuria_proband, prader_willi_proband, rett_syndrome_proband, other_genetic_proband, smith_lemli_opitz_proband |
| 14 | any_neurological_disorders_proband | OR | other_neurological_disorder_proband, cerebral_palsy_proband, congenital_rubella_proband, cranial_nerve_disorder_proband, hydrocephalus_proband, landau_kleffner_syndrome_proband, neurofibromatosis_proband, tuberous_sclerosis_proband, migraines_proband, seizures_proband |
| 15 | any_neurological_disorders_in_family | OR | other_neurological_disorder_y_n_in_family, cerebral_palsy_y_n_in_family, congenital_rubella_y_n_in_family, cranial_nerve_disorder_y_n_in_family, hydrocephalus_y_n_in_proband, landau_kleffner_syndrome_y_n_in_family, neurofibromatosis_y_n_in_family, tuberous_sclerosis_y_n_in_family, migraines_y_n_in_family, seizures_y_n_in_family |
| 16 | count_of_immunizations_of_proband | COUNT "TRUE" | child_chicken_pox_varicella, child_dpt, child_dtap, child_flu_shot, child_hib, child_mmr, child_hepatitis_b, child_polio_injected, child_polio_oral, child_vaccination_other_1_y_n, child_vaccination_other_2_y_n, child_tetanus_booster, child_travel_related_vaccines |
| 17 | child_diet_casein_past_or_present | OR | child_diet_casein_current, child_diet_casein_past |
| 18 | child_vitamins_past_or_present | OR | child_vitamins, child_vitamins_past |
| 19 | child_diet_gluten_past_or_present | OR | child_diet_gluten_current, child_diet_gluten_past |
| 20 | child_diets_helpful_past_or_present | OR | child_diet_helpful_current, child_diet_helpful_past |
| 21 | mother_age_at_conception | SUBTRACT | proband_year_of_birth, mother_year_of_birth |
| 22 | father_age_at_conception | SUBTRACT | proband_year_of_birth, father_year_of_birth |
| 23 | avg_temperature_ados_day | AVERAGE | temperature1, temperature2, temperature3 |
| 24 | years_treated_at_home | COUNT ("Home"\| "Both") | age02_intensive_behav_therapy_prog_typ, …, age17_intensive_behav_therapy_prog_typ |
| 25 | ever_treated_at_home_intesive _therapy | OR ("Home"\| "Both") | age02_intensive_behav_therapy_prog_typ, …, age17_intensive_behav_therapy_prog_typ |
| 26 | any_maternal_autoimmune_disease | OR | allergies_trimester_1, allergies_trimester_2, allergies_trimester_3, adrenal_insufficiency_mat, bowel_disorders_mat, celiac_disease_mat, diabetes_mellitus_type1_mat, hashimotos_tyroiditis_mat, hyperthyroidism_mat, hypothyroidism_mat, multiple_sclerosis_mat, other_autoimmune_disorder_mat, psoriasis_mat, rheum_arthritis_adult_mat, rheum_arthritis_juvenile_mat, systemic_lupus_eryth_mat |
| 27 | any_maternal_infection_in _pregnancy | OR | medhx_preg_illneses_substances.chicken_pox, medhx_preg_illneses_substances.enterovirus,  medhx_preg_illneses_substances.herpes_1_type_1, medhx_preg_illneses_substances.herpes_2_type_2, medhx_preg_illneses_substances.infectious_mono, medhx_preg_illneses_substances.influenza,  medhx_preg_illneses_substances.kidney_infection, medhx_preg_illneses_substances.mat_fever,  medhx_preg_illneses_substances.vaginal_infection, medhx_preg_illneses_substances.viral_hepatitis, medhx_preg_illneses_substances.other_viral_illness_1, medhx_preg_illneses_substances.other_viral_illness_2, medhx_preg_illneses_substances.uti, medhx_preg_illneses_substances.skin_rash, medhx_preg_illneses_substances.upper_respiratory_virus |
| 28 | celiac_family_excluding_proband | In family and not in proband | cellac_disease_y_n, cellac_disease_proband |
| 29 | bowel_disorders_family_excluding_ proband | In family and not in proband | bowel_disorders_proband, bowel_disorders_y_n |

**Supplementary Table 3** Continuous variable clusters and their relation to fever response

| Cluster Number | Feature | Responders | | Non-Responders | | Cluster P | Cluster FDR |
| --- | --- | --- | --- | --- | --- | --- | --- |
|  |  | Mean | SEM | Mean | SEM |  |  |
| 1 | adi_r_b_comm_verbal_total | 17.9 | 0.15 | 16.23 | 0.15 | < 1x10-6 | < 1x10-6 |
|  | adi_r_comm_b_non_verbal_total | 10.23 | 0.12 | 9.12 | 0.13 |  |  |
| 2 | verbal_iq | 67.1 | 1.19 | 78.78 | 1.13 | < 1x10-6 | < 1x10-6 |
|  | nonverbal_iq | 76.01 | 0.99 | 84.57 | 0.94 |  |  |
|  | vabs_ii_soc_standard | 67.46 | 0.46 | 71.36 | 0.46 |  |  |
|  | vabs_ii_communication | 73.41 | 0.55 | 77.29 | 0.53 |  |  |
|  | vabs_ii_dls_standard | 73.52 | 0.52 | 76.3 | 0.49 |  |  |
| 3 | adi_r_soc_a_total | 22.1 | 0.19 | 20.03 | 0.21 | 1.00E-06 | 1.40E-05 |
| 4 | abc_iii_stereotypy | 6.29 | 0.18 | 4.68 | 0.15 | 4.00E-06 | 4.20E-05 |
| 5 | adi_r_rrb_c_total | 7.14 | 0.09 | 6.33 | 0.09 | 1.10E-05 | 9.24E-05 |
| 6 | abc_total_score | 53.37 | 1 | 45.1 | 0.93 | 1.70E-05 | 0.000119 |
|  | abc_iv_hyperactivity | 19.14 | 0.4 | 16.17 | 0.38 |  |  |
|  | cbcl_6_18_thought_problems | 69.14 | 0.3 | 66.62 | 0.31 |  |  |
|  | abc_i_irritability | 12.95 | 0.34 | 11.13 | 0.31 |  |  |
|  | cbcl_6_18_aggressive_behavior | 60.77 | 0.35 | 59.02 | 0.32 |  |  |
|  | cbcl_6_18_externalizing_t_score | 57.2 | 0.39 | 55.64 | 0.38 |  |  |
|  | cbcl_6_18_oppositional_defiant | 59.82 | 0.32 | 58.45 | 0.29 |  |  |
|  | cbcl_6_18_total_problems | 63.44 | 0.31 | 62.26 | 0.31 |  |  |
|  | cbcl_6_18_conduct_problems | 57.74 | 0.29 | 56.82 | 0.27 |  |  |
|  | cbcl_6_18_anxiety_problems | 61.9 | 0.31 | 61.23 | 0.32 |  |  |
|  | cbcl_6_18_social_problems | 62.52 | 0.29 | 63.11 | 0.29 |  |  |
|  | cbcl_6_18_affective_problems | 61.89 | 0.34 | 61.51 | 0.31 |  |  |
|  | cbcl_6_18_rule_breaking | 55.69 | 0.24 | 55.37 | 0.22 |  |  |
|  | cbcl_6_18_anxious_depressed | 58.73 | 0.32 | 59.16 | 0.33 |  |  |
|  | cbcl_6_18_withdrawn | 62.89 | 0.31 | 62.79 | 0.32 |  |  |
|  | cbcl_6_18_internalizing_t_score | 59.73 | 0.34 | 59.58 | 0.36 |  |  |
| 7 | srs_parent_t_score | 81.7 | 0.35 | 78.95 | 0.39 | 0.000285 | 0.00171 |
| 8 | rbs_r_overall_score | 30.73 | 0.67 | 26.35 | 0.63 | 0.000506 | 0.002654 |
| 9 | sum_severity_gastro_disorders | 2.91 | 0.14 | 2 | 0.11 | 0.001069 | 0.004834 |
|  | max_severity_gastro_disorders | 1.47 | 0.05 | 1.2 | 0.05 |  |  |
| 10 | abc_v_inappropriate_speech | 4.27 | 0.12 | 3.47 | 0.11 | 0.001151 | 0.004834 |
| 11 | diarrhea_specify | 0.45 | 0.04 | 0.25 | 0.03 | 0.002566 | 0.009797 |
| 12 | SRS_social_cognition_mother | 5.37 | 0.17 | 6.37 | 0.19 | 0.00394 | 0.01379 |
|  | SRS_total_mother | 26.3 | 0.66 | 30.33 | 0.76 |  |  |
|  | SRS_communication_mother | 7.42 | 0.22 | 8.69 | 0.27 |  |  |
|  | SRS_social_awareness_mother | 4.19 | 0.1 | 4.58 | 0.1 |  |  |
|  | SRS_mannerisms_mother | 3.92 | 0.13 | 4.56 | 0.15 |  |  |
|  | SRS_motivation_mother | 5.4 | 0.16 | 6.14 | 0.18 |  |  |
| 13 | cbcl_6_18_social | 32.33 | 0.33 | 34.59 | 0.34 | 0.007388 | 0.023867 |
| 14 | total_DCDQ | 36.45 | 0.43 | 38.86 | 0.46 | 0.012528 | 0.037584 |
|  | control_during_movement | 15.53 | 0.2 | 16.59 | 0.22 |  |  |
|  | fine_motor_handwriting | 9.37 | 0.16 | 10.13 | 0.17 |  |  |
|  | general_coordination | 11.6 | 0.15 | 12.13 | 0.16 |  |  |
| 15 | bloating_specify | 0.21 | 0.03 | 0.1 | 0.02 | 0.015451 | 0.043263 |
| 16 | abc_ii_lethargy | 10.71 | 0.26 | 9.65 | 0.26 | 0.016946 | 0.044482 |
| 17 | cbcl_6_18_attention_problems | 68.54 | 0.38 | 66.49 | 0.37 | 0.018069 | 0.04464 |
|  | cbcl_6_18_add_adhd | 62.92 | 0.3 | 61.53 | 0.29 |  |  |
| 18 | phrase_delay | 0.81 | 0.02 | 0.7 | 0.02 | 0.030585 | 0.071365 |
| 19 | mother_age_at_conception | 30.89 | 0.18 | 31.52 | 0.18 | 0.071098 | 0.152413 |
|  | father_age_at_conception | 33.3 | 0.22 | 33.66 | 0.21 |  |  |
| 20 | standard_score_CTOPP | 7.21 | 0.1 | 7.73 | 0.1 | 0.074901 | 0.152413 |
| 21 | number_of_distinct_autoimmune  _diseases_in_family | 2.08 | 0.05 | 1.88 | 0.05 | 0.076207 | 0.152413 |
| 22 | bowel_disorders_family_excluding_proband | 0.14 | 0.01 | 0.1 | 0.01 | 0.096944 | 0.181661 |
| 23 | excessive_gas_specify | 0.27 | 0.03 | 0.17 | 0.02 | 0.099481 | 0.181661 |
| 24 | abc_nbr_missing | 0.01 | 0 | 0.09 | 0.05 | 0.156469 | 0.273821 |
| 25 | SRS_social_cognition_father | 4.47 | 0.16 | 4.91 | 0.17 | 0.16772 | 0.281769 |
|  | SRS_social_awareness_father | 4.8 | 0.12 | 4.96 | 0.11 |  |  |
|  | SRS_total_father | 28.38 | 0.82 | 29.69 | 0.82 |  |  |
|  | SRS_mannerisms_father | 3.68 | 0.15 | 3.94 | 0.16 |  |  |
|  | SRS_communication_father | 9.35 | 0.31 | 9.61 | 0.3 |  |  |
|  | SRS_motivation_father | 6.08 | 0.2 | 6.26 | 0.2 |  |  |
| 26 | celiac_family_excluding_proband | 0.04 | 0.01 | 0.02 | 0.01 | 0.194683 | 0.307367 |
| 27 | constipation_specify | 0.75 | 0.04 | 0.64 | 0.04 | 0.204911 | 0.307367 |
| 28 | unusual_stools_specify | 0.45 | 0.04 | 0.36 | 0.03 | 0.198855 | 0.307367 |
| 29 | ados_css | 4.54 | 0.07 | 4.46 | 0.07 | 0.284039 | 0.40703 |
| 30 | non_febrile_seizures | 0.2 | 0.02 | 0.15 | 0.02 | 0.290736 | 0.40703 |
| 31 | vomiting_specify | 0.14 | 0.02 | 0.11 | 0.02 | 0.330073 | 0.447196 |
| 32 | height_z_score | 0.33 | 0.04 | 0.39 | 0.04 | 0.437526 | 0.574253 |
| 33 | head_circumference | 53.88 | 0.09 | 53.96 | 0.1 | 0.536579 | 0.666375 |
|  | height | 133.2 | 0.73 | 134.27 | 0.78 |  |  |
|  | weight | 35.17 | 0.67 | 36.05 | 0.73 |  |  |
|  | age_at_ados | 8.82 | 0.13 | 8.9 | 0.13 |  |  |
| 34 | cbcl_6_18_somatic_prob | 55.94 | 0.29 | 55.43 | 0.27 | 0.559143 | 0.666375 |
|  | cbcl_6_18_somatic_complaints | 57.06 | 0.29 | 56.68 | 0.27 |  |  |
| 35 | num_sibs | 1.38 | 0.03 | 1.37 | 0.03 | 0.666375 | 0.666375 |
| 36 | febrile_seizures | 0.06 | 0.01 | 0.06 | 0.01 | 0.571961 | 0.666375 |
| 37 | word_delay | 0.42 | 0.02 | 0.4 | 0.02 | 0.571262 | 0.666375 |
| 38 | count_of_immunizations_of_proband | 6.58 | 0.06 | 6.56 | 0.06 | 0.66259 | 0.666375 |
| 39 | proband_birth_order | 1.75 | 0.03 | 1.72 | 0.03 | 0.627702 | 0.666375 |
| 40 | annual_household_income | 5.86 | 0.08 | 5.84 | 0.08 | 0.66567 | 0.666375 |
| 41 | bmi_z_score | 0.68 | 0.05 | 0.67 | 0.05 | 0.641828 | 0.666375 |
| 42 | cbcl_6_18_activities | 39.05 | 0.39 | 39.26 | 0.35 | 0.633077 | 0.666375 |

**Supplementary Table 4** Categorical variable clusters and their association with fever response

| Cluster Number | Feature | OR [95% CI] | Cluster P | Cluster FDR |
| --- | --- | --- | --- | --- |
| 1 | child_diet_gluten_past_or_present | 2.42 [1.97, 2.99] | 2.00E-06 | 0.000206 |
|  | child_vitamins_past_or_present | 2.13 [1.74, 2.59] |  |  |
|  | child_diet_casein_past_or_present | 2.12 [1.7, 2.65] |  |  |
|  | food_allergies_milk | 2.2 [1.68, 2.91] |  |  |
|  | food_allergies_casein | 3.1 [1.96, 4.96] |  |  |
|  | any_gastro_disorders_proband | 1.46 [1.15, 1.81] |  |  |
|  | food_allergies_gluten | 2.28 [1.43, 3.52] |  |  |
|  | environment_allergies_pollen | 1.27 [0.97, 1.66] |  |  |
|  | food_allergies_nuts | 1.48 [0.99, 2.21] |  |  |
|  | food_allergies_egg | 1.53 [0.94, 2.39] |  |  |
|  | food_allergies_wheat | 1.5 [0.91, 2.43] |  |  |
|  | environment_allergies_dust | 0.87 [0.57, 1.27] |  |  |
|  | food_allergies_fish_shellfish | 1.32 [0.87, 2.24] |  |  |
| 2 | nonstandardized_impressions | NA | 0.000279 | 0.0143685 |
| 3 | any_maternal_infections_in_preg | 1.7 [1.42, 2.03] | 0.000424 | 0.014557333 |
|  | upper_respiratory_virus | 1.3 [1.11, 1.53] |  |  |
|  | mat_fever | 1.75 [1.17, 3.06] |  |  |
|  | enterovirus | 1.58 [1.08, 2.37] |  |  |
|  | uti | 1.32 [1.01, 1.79] |  |  |
|  | influenza | 1.28 [0.89, 2.04] |  |  |
|  | other_viral_illness_1 | 1.18 [0.78, 1.7] |  |  |
|  | herpes_2_type_2 | 0.9 [0.43, 2.94] |  |  |
|  | herpes_1_type_1 | 1 [0.3, 2.1] |  |  |
|  | skin_rash | 1 [0.64, 1.65] |  |  |
|  | vaginal_infection | 1.07 [0.7, 1.69] |  |  |
|  | other_viral_illness_2 | 0 [0, 0] |  |  |
| 4 | food_allergies | NA | 0.002979 | 0.07670925 |
| 5 | diarrhea_y_n | 1.87 [1.45, 2.59] | 0.004502 | 0.0927412 |
| 6 | severe_abdominal_pain_y_n | NA | 0.013952 | 0.239509333 |
| 7 | cpea_dx | NA | 0.026443 | 0.376194625 |
| 8 | communication_disorder_proband | 3.37 [2, 6.11] | 0.029219 | 0.376194625 |
|  | receptive_lang_disorder_proband | 2.6 [1.6, 4.53] |  |  |
|  | expressive_lang_disorder_proband | 1.99 [1.36, 2.94] |  |  |
|  | communication_disorder_y_n_in_family | 2.29 [1.19, 4.13] |  |  |
|  | receptive_lang_disorder_y_n_in_family | 2.07 [1.13, 3.97] |  |  |
|  | mixed_expressive_disorder_proband | 0.67 [0.43, 0.97] |  |  |
|  | expressive_lang_disorder_y_n_in_family | 1.51 [0.95, 2.34] |  |  |
|  | speech_delay_proband | 0.81 [0.65, 1] |  |  |
|  | mixed_expressive_disorder_y_n_in_family | 0.72 [0.51, 0.98] |  |  |
|  | nonverbal_disorder_y_n | 1.92 [1.01, 4.16] |  |  |
|  | stuttering_y_n_in_family | 1.32 [0.92, 1.84] |  |  |
|  | pervasive_developmental_nos_y_n | 0.8 [0.61, 1.03] |  |  |
|  | reading_disorder_y_n | 1.25 [0.94, 1.68] |  |  |
|  | any_language_disorders_proband | 0.9 [0.75, 1.08] |  |  |
|  | pragmatics_lang_disorder_proband | 0.73 [0.46, 1.12] |  |  |
|  | written_expression_disorder_y_n | 0.68 [0.19, 1.45] |  |  |
|  | speech_delay_y_n_in_family | 0.92 [0.73, 1.14] |  |  |
|  | reduced_articulation_y_n_in_family | 1.09 [0.85, 1.38] |  |  |
|  | stuttering_proband | 1.32 [0.4, 3.26] |  |  |
|  | any_language_disorders_in_family | 0.96 [0.79, 1.17] |  |  |
|  | reduced_articulation_proband | 1.04 [0.68, 1.55] |  |  |
|  | pragmatics_lang_disorder_y_n_in_family | 0.94 [0.66, 1.33] |  |  |
|  | math_disorder_y_n | 1.2 [0.66, 2.1] |  |  |
| 9 | systemic_lupus_eryth_y_n_in_family | 2.08 [1.52, 2.91] | 0.034791 | 0.3877847 |
|  | hypothyroidism_proband | 6.01 [2.87, 16.15] |  |  |
|  | psoriasis_proband | 2.94 [1.87, 5.03] |  |  |
|  | cellac_disease_y_n_in_family | 2.03 [1.14, 3.56] |  |  |
|  | any_autoimmune_disease_in_family | 1.36 [1.03, 1.72] |  |  |
|  | cellac_disease_proband | 4.12 [1.15, 12.79] |  |  |
|  | asthma_y_n_in_family | 1.22 [0.98, 1.51] |  |  |
|  | hashimotos_tyroiditis_y_n_in_family | 1.61 [1.01, 2.64] |  |  |
|  | bowel_disorders_proband | 1.7 [0.95, 2.95] |  |  |
|  | asthma_mat | 1.25 [0.9, 1.73] |  |  |
|  | hyperthyroidism_y_n_in_family | 1.23 [0.79, 1.86] |  |  |
|  | severe_allergies | 0.65 [0.37, 1.3] |  |  |
|  | any_autoimmune_disease_mat | 1.05 [0.85, 1.33] |  |  |
|  | any_autoimmune_disease_proband | 0.93 [0.73, 1.25] |  |  |
|  | hypothyroidism_y_n_in_family | 1.03 [0.8, 1.33] |  |  |
|  | asthma_proband | 0.91 [0.6, 1.34] |  |  |
|  | psoriasis_y_n_in_family | 0.97 [0.68, 1.35] |  |  |
| 10 | bloating_y_n | NA | 0.037649 | 0.3877847 |
| 11 | bowel_disorders_y_n_in_family | 1.67 [1.29, 2.17] | 0.046027 | 0.430980091 |
|  | asperger_syndrome_y_n | 0.55 [0.4, 0.75] |  |  |
|  | anxiety_disorder_y_n_in_family | 1.24 [1, 1.53] |  |  |
|  | admitted_hospital_psych_proband | NA |  |  |
|  | anxiety_disorder_proband | 0.5 [0.29, 0.93] |  |  |
|  | personality_disorder_y_n_in_family | 1.78 [1.03, 3.57] |  |  |
|  | eating_disorder_y_n_in_family | 1.28 [0.85, 1.87] |  |  |
|  | attention_deficit_proband | 0.8 [0.58, 1.11] |  |  |
|  | depressive_disorder_proband | 0 [0, 0] |  |  |
|  | bipolar_disorder_y_n_in_family | 0.86 [0.62, 1.17] |  |  |
|  | social_phobia_y_n_in_family | 1.21 [0.71, 2.04] |  |  |
|  | depressive_disorder_y_n_in_family | 0.92 [0.74, 1.15] |  |  |
|  | obsessive_compulsive_y_n_in_family | 0.89 [0.58, 1.3] |  |  |
|  | post_traumatic_stress_y_n_in_family | 1.19 [0.73, 1.88] |  |  |
|  | schizophrenia_y_n_in_family | 0.86 [0.5, 1.41] |  |  |
|  | behavior_disorder_proband | 0.7 [0, 2.21] |  |  |
|  | attention_deficit_y_n_in_family | 1.05 [0.84, 1.3] |  |  |
|  | pica_proband | 1.12 [0.33, 2.76] |  |  |
|  | pica_y_n_in_family | 1.2 [0.53, 2.51] |  |  |
|  | obsessive_compulsive_proband | 0.99 [0.45, 2] |  |  |
|  | admitted_hospital_psych_y_n_in_family | 1.01 [0.75, 1.35] |  |  |
|  | behavior_disorder_y_n_in_family | 0.95 [0.42, 1.81] |  |  |
|  | eating_disorder_proband | 1.26 [0, 4.24] |  |  |
|  | bipolar_disorder_proband | 0.99 [0, 4.17] |  |  |
|  | social_phobia_proband | NA |  |  |
|  | post_traumatic_stress_proband | 0 [0, 0] |  |  |
| 12 | pos_lead_y_n | NA | 0.072669 | 0.534636214 |
| 13 | strep_throat_y_n | NA | 0.069092 | 0.534636214 |
| 14 | gastroesophageal_reflux_y_n | NA | 0.070868 | 0.534636214 |
| 15 | other_disorder_illness1_y_n_in_family | 1.42 [1.14, 1.76] | 0.088464 | 0.573291941 |
|  | other_autoimmune_disorder_y_n_in_family | 1.51 [1.14, 2] |  |  |
|  | other_disorder_illness2_y_n_in_family | 1.17 [0.68, 1.9] |  |  |
|  | child_vaccination_other_1_y_n | 1.09 [0.74, 1.52] |  |  |
|  | child_vaccination_other_2_y_n | 0.95 [0.51, 1.51] |  |  |
|  | rheum_arthritis_adult_y_n_in_family | 0.96 [0.71, 1.26] |  |  |
|  | diabetes_mellitus_type1_y_n_in_family | 0.9 [0.56, 1.38] |  |  |
| 16 | any_chronic_illnesses_in_family | 1.45 [1.13, 1.83] | 0.094621 | 0.573291941 |
|  | cancer_y_n_in_family | 1.2 [0.96, 1.5] |  |  |
|  | diabetes_mellitus_type2_y_n_in_family | 1.09 [0.87, 1.36] |  |  |
|  | death_under_50_y_n_in_family | 1 [0.7, 1.39] |  |  |
|  | heart_disease_y_n_in_family | 0.97 [0.79, 1.2] |  |  |
|  | stroke_y_n_in_family | 0.95 [0.71, 1.25] |  |  |
|  | diabetes_mellitus_type2_mat | 0.87 [0.33, 2.15] |  |  |
| 17 | excessive_gas_y_n | NA | 0.089152 | 0.573291941 |
| 18 | adjustment_disorder_y_n_in_family | 3.55 [1.38, 11.85] | 0.129281 | 0.739774611 |
|  | adjustment_disorder_proband | 3.84 [0, 8.99] |  |  |
| 19 | celiac_disease_gas_ill_y_n | NA | 0.16752 | 0.838012905 |
| 20 | head_injury_y_n | NA | 0.170857 | 0.838012905 |
| 21 | metabolic_syn_y_n | NA | 0.167109 | 0.838012905 |
| 22 | fragile_x_y_n_in_family | NA | 0.19214 | 0.886592652 |
| 23 | hydrocephalus_proband | NA | 0.197977 | 0.886592652 |
| 24 | migraines_proband | 2.43 [1.07, 5.94] | 0.212535 | 0.912129375 |
|  | other_neurological_disorder_proband | NA |  |  |
|  | migraines_y_n_in_family | 1.22 [0.98, 1.5] |  |  |
|  | any_neurological_disorders_proband | 1.36 [1, 2.09] |  |  |
|  | any_neurological_disorders_in_family | 1.16 [0.97, 1.4] |  |  |
|  | other_neurological_disorder_y_n_in_family | 1.4 [0.93, 2.11] |  |  |
|  | cranial_nerve_disorder_y_n_in_family | 1.49 [0.8, 2.68] |  |  |
|  | hydrocephalus_y_n_in_family | 0.9 [0, 4.87] |  |  |
|  | seizures_y_n_in_family | 1.06 [0.8, 1.37] |  |  |
|  | seizures_proband | 0.92 [0.54, 1.48] |  |  |
| 25 | cleft_lip_palate_y_n_in_family | 1.87 [1.16, 3.35] | 0.29467 | 1 |
|  | any_birth_defects_proband | 1.4 [0.9, 2.65] |  |  |
|  | cleft_lip_palate_proband | 2.04 [0, 6.42] |  |  |
|  | abnormal_shape_polydactyly_proband | 0 [0, 1.04] |  |  |
|  | any_birth_defects_in_family | 1.06 [0.86, 1.35] |  |  |
|  | open_spine_y_n_in_family | 0.58 [0, 1.96] |  |  |
|  | other_birth_defect_proband | 0.64 [0, 2.17] |  |  |
|  | other_birth_defect_y_n_in_family | 1.11 [0.73, 1.61] |  |  |
|  | congenital_heart_defect_y_n_in_family | 1.05 [0.72, 1.5] |  |  |
|  | abnormal_shape_polydactyly_y_n_in_family | 0.74 [0.21, 1.54] |  |  |
|  | kidney_defect_y_n_in_family | 0.86 [0.41, 1.77] |  |  |
|  | congenital_heart_defect_proband | 0 [0, 1.93] |  |  |
|  | kidney_defect_proband | 0 [0, 2.73] |  |  |
| 26 | child_mmr | 0.49 [0.25, 0.9] | 0.284897 | 1 |
|  | child_hib | 0.65 [0.4, 1.06] |  |  |
|  | child_dpt | 1.31 [0.96, 1.78] |  |  |
|  | child_chicken_pox_varicella | 0.81 [0.62, 1.07] |  |  |
|  | child_hepatitis_b | 0.78 [0.48, 1.27] |  |  |
|  | child_tetanus_booster | 0.88 [0.68, 1.12] |  |  |
|  | vision | 0.9 [0.65, 1.21] |  |  |
|  | child_polio_oral | 0.89 [0.62, 1.25] |  |  |
|  | child_flu_shot | 0.98 [0.78, 1.24] |  |  |
|  | child_dtap | 0.91 [0.65, 1.24] |  |  |
|  | child_polio_injected | 1.01 [0.78, 1.29] |  |  |
| 27 | mental_retardation_y_n | 1.44 [0.94, 2.19] | 0.324717 | 1 |
|  | other_genetic_y_n_in_family | 1.41 [0.93, 2.19] |  |  |
|  | down_syndrome_y_n_in_family | 1.68 [0.72, 3.99] |  |  |
|  | other_genetic_proband | 0.38 [0, 1.28] |  |  |
|  | any_genetic_disorders_in_family | 1.17 [0.84, 1.76] |  |  |
|  | any_genetic_disorders_proband | 0.75 [0.42, 3.01] |  |  |
| 28 | adrenal_insufficiency_mat | 0 [0, 4.34] | 1 | 1 |
| 29 | adrenal_insufficiency_y_n_in_family | 0 [0, 2.09] | 1 | 1 |
| 30 | landau_kleffner_syndrome_proband | 0 [0, 0] | 1 | 1 |
| 31 | landau_kleffner_syndrome_y_n_in_family | 0 [0, 0] | 1 | 1 |
| 32 | cerebral_palsy_y_n_in_family | 0.29 [0, 0.95] | 0.32243 | 1 |
|  | cerebral_palsy_proband | 0 [0, 0] |  |  |
| 33 | dysthymic_disorder_y_n_in_family | 0.42 [0.22, 0.99] | 0.486132 | 1 |
|  | dysthymic_disorder_proband | 0 [0, 2] |  |  |
| 34 | tourettes_proband | 0 [0, 0.87] | 0.589083 | 1 |
|  | tourettes_y_n_in_family | 0.6 [0.31, 1.13] |  |  |
| 35 | irritable_bowel_y_n | NA | 0.793443 | 1 |
| 36 | race | NA | 0.568894 | 1 |
| 37 | family_type | 0.82 [0.67, 1.04] | 0.347673 | 1 |
| 38 | sex | 1.1 [0.81, 1.37] | 0.669039 | 1 |
| 39 | vomiting_y_n | NA | 0.42756 | 1 |
| 40 | crohns_disease_y_n | 0 [0, 0] | 1 | 1 |
| 41 | inflammatory_bowel_y_n | NA | 0.561419 | 1 |
| 42 | child_travel_related_vaccines | 0.77 [0.39, 1.45] | 0.779295 | 1 |
| 43 | diabetes_mellitus_type1_proband | NA | 1 | 1 |
| 44 | diabetes_mellitus_type2_proband | NA | 1 | 1 |
| 45 | hashimotos_tyroiditis_proband | 0 [0, 0] | 1 | 1 |
| 46 | hyperthyroidism_proband | NA | 1 | 1 |
| 47 | multiple_sclerosis_y_n_in_family | 0.8 [0.45, 1.37] | 0.799914 | 1 |
| 48 | multiple_sclerosis_proband | 0 [0, 0] | 1 | 1 |
| 49 | rheum_arthritis_adult_proband | NA | 1 | 1 |
| 50 | rheum_arthritis_juvenile_y_n_in_family | 1.65 [1, 2.81] | 0.354438 | 1 |
| 51 | rheum_arthritis_juvenile_proband | 0 [0, 0] | 1 | 1 |
| 52 | systemic_lupus_eryth_proband | 0 [0, 0] | 1 | 1 |
| 53 | open_spine_proband | NA | 1 | 1 |
| 54 | angelman_y_n_in_family | 0 [0, 0] | 1 | 1 |
| 55 | phenylketonuria_y_n_in_family | 0 [0, 0] | 1 | 1 |
| 56 | phenylketonuria_proband | 0 [0, 0] | 1 | 1 |
| 57 | prader_willi_y_n_in_family | 0 [0, 0] | 1 | 1 |
| 58 | childhood_disintegration_y_n | NA | 1 | 1 |
| 59 | other_psychotic_disorder_y_n_in_family | 1.44 [0.69, 4.53] | 0.57377 | 1 |
| 60 | cerebral_palsy_neuro_probs_y_n | NA | 0.749335 | 1 |
| 61 | encephalitis_y_n | NA | 1 | 1 |
| 62 | excess_clumsy_uncoordinated_yn | NA | 0.418723 | 1 |
| 63 | febrile_seizures_y_n | NA | 0.766986 | 1 |
| 64 | meningitis_y_n | 1.76 [0.88, 3.76] | 0.51459 | 1 |
| 65 | movement_abnormalities_y_n | NA | 0.726607 | 1 |
| 66 | tuberous_sclerosis_neuro_y_n | 0 [0, 0] | 1 | 1 |
| 67 | tourettes_tics_other_y_n | NA | 0.785477 | 1 |
| 68 | bckgd_hx_parent_relation_status | NA | 0.902723 | 1 |
| 69 | family_structure | NA | 0.507584 | 1 |
| 70 | chicken_pox_infect_disease_y_n | NA | 0.801378 | 1 |
| 71 | epstein_barr_virus_y_n | NA | 1 | 1 |
| 72 | genital_probs_y_n | 0.58 [0.33, 1.35] | 0.529664 | 1 |
| 73 | heart_probs_y_n | NA | 0.440563 | 1 |
| 74 | herpes_1_y_n | 1.26 [0.83, 2.1] | 0.685186 | 1 |
| 75 | kidney_urinary_probs_y_n | 1.26 [0.76, 3.39] | 0.672834 | 1 |
| 76 | lyme_disease_y_n | 0 [0, 0.97] | 0.55476 | 1 |
| 77 | measles_infect_disease_y_n | 0 [0, 0] | 1 | 1 |
| 78 | mumps_infect_disease_y_n | 0 [0, 0] | 1 | 1 |
| 79 | otitis_media_y_n | NA | 0.341872 | 1 |
| 80 | respiratory_probs_y_n | NA | 0.778484 | 1 |
| 81 | roseola_y_n | NA | 0.780864 | 1 |
| 82 | rubella_infect_disease_y_n | NA | 1 | 1 |
| 83 | surgeries_y_n | NA | 0.506297 | 1 |
| 84 | highest_edu_father | NA | 0.899361 | 1 |
| 85 | highest_edu_mother | NA | 0.853855 | 1 |
| 86 | parent_relation_status | NA | 0.863965 | 1 |
| 87 | chicken_pox | 0 [0, 0] | 1 | 1 |
| 88 | infectious_mono | 0 [0, 0] | 1 | 1 |
| 89 | kidney_infection | 0 [0, 0] | 1 | 1 |
| 90 | viral_hepatitis | NA | 1 | 1 |
| 91 | constipation_y_n | NA | 0.389656 | 1 |
| 92 | gastric_ulcera_y_n | NA | 1 | 1 |
| 93 | unusual_stools_y_n | 1 [0.8, 1.25] | 0.821659 | 1 |
| 94 | ulcerative_colitis_y_n | 0 [0, 0] | 1 | 1 |
| 95 | color_blind | NA | 0.570714 | 1 |
| 96 | environment_allergies | NA | 0.638811 | 1 |
| 97 | environment_allergies_latex | 2.15 [0.9, 8.47] | 0.465013 | 1 |
| 98 | hearing | 1.31 [0.88, 2.04] | 0.522721 | 1 |
| 99 | handedness | NA | 0.651816 | 1 |
| 100 | medications_allergies | NA | 0.597746 | 1 |
| 101 | congenital_rubella_y_n_in_family | 0 [0, 0] | 1 | 1 |
| 102 | neurofibromatosis_y_n_in_family | 0 [0, 4.25] | 1 | 1 |
| 103 | neurofibromatosis_proband | NA | 1 | 1 |
| 104 | tuberous_sclerosis_y_n_in_family | 0 [0, 0] | 1 | 1 |
| 105 | tuberous_sclerosis_proband | 0 [0, 0] | 1 | 1 |
| 106 | is_hispanic | 1.11 [0.85, 1.41] | 0.649895 | 1 |
